# Evidence for rapid evolution in a grassland biodiversity experiment

**DOI:** 10.1101/262303

**Authors:** Sofia J. van Moorsel, Marc W. Schmid, Niels C.A.M. Wagemaker, Thomas van Gurp, Bernhard Schmid, Philippine Vergeer

## Abstract

Biodiversity often increases plant productivity. In long-term grassland experiments, positive biodiversity effects on plant productivity commonly increase with time. Also, it has been shown that such positive biodiversity effects persist not only in the local environment but also when plants are transferred into a common environment. Thus, we hypothesized that community diversity had acted as a selective agent, resulting in the emergence of plant monoculture and mixture types with differing genetic composition. To test our hypothesis, we grew offspring from plants that were grown for eleven years in monoculture or mixture environments in a biodiversity experiment (Jena Experiment) under controlled glasshouse conditions in monocultures or two-species mixtures. We used epiGBS, a genotyping-by-sequencing approach combined with bisulfite conversion to provide integrative genetic and epigenetic data. We observed significant genetic and epigenetic divergence according to selection history in three out of five perennial grassland species, namely *Galium mollugo*, *Prunella vulgaris* and *Veronica chamaedrys*, with epigenetic differences mostly reflecting the genetic differences. In addition, current diversity levels in the glasshouse had weak effects on epigenetic variation. However, given the limited genome coverage of the reference-free bisulfite method epiGBS, it remains unclear how much of this epigenetic divergence was independent of underlying genetic differences. Our results thus suggest that selection of genetic variants, and possibly epigenetic variants, caused the rapid emergence of monoculture and mixture types within plant species in the Jena Experiment.

## 1. Introduction

Environmental change may result in significant climate-induced range shifts, forcing plant populations into new abiotic and biotic environments (Ouborg, Vergeer, & Mix 2006). The unprecedented rate of environmental change raises the question whether adaptation of natural communities to novel abiotic or biotic conditions is able to match these rates. Biodiversity is known to buffer ecosystems against negative influences of climatic extremes and novel environmental conditions (Isbell *et al*., 2015) and in addition it has been shown that co-evolution among plants comprising a community can dampen the impact of an extreme climatic event (van Moorsel *et al*. 2018a).

Adaptive responses of plant populations to environmental conditions (e.g., Joshi *et al*. 2001) and biotic interactions such as pollinators (e.g., Gervasi & Schiestl 2017) are well studied. Unfortunately, little effort has been devoted to the influence of community diversity on fast evolutionary adaptation (but see Lipowsky, Schmid, & Roscher 2011, Kleynhans, Otto, Reich, & Vellend 2016, van Moorsel *et al*. 2018b). In particular, potential effects of multispecies interactions on the adaptive response of plant species are largely unknown, despite a growing body of evidence showing the importance of multispecies interactions for ecosystem stability (Bastolla *et al*. 2009, Guimarães, Pires, Jordano, Bascompte, & Thompson 2017). It is conceivable that the feedback between species interactions and their adaptive responses shapes community-level ecosystem functioning (van Moorsel *et al*. 2018b).

In the 1960s it was proposed that evolutionary processes occur at longer time scales than ecological processes (Slobodkin 1961), but now it is commonly believed that micro-evolutionary and ecological processes can occur at the same or at least at similar temporal scales (Hairston, Ellner, Geber, Yoshida, & Fox 2005, Schoener 2011, Hendry 2016). A good understanding of how biodiversity, i.e. the interaction between species, shapes evolutionary responses, is instrumental for predicting ecosystem responses to global change and biodiversity loss.

Long-term biodiversity field experiments offer unique opportunities to study effects of community diversity and composition on natural selection. Species mixtures are frequently more productive than average monocultures (Balvanera *et al*. 2006). Interestingly, these biodiversity effects often become more pronounced over time, which has been attributed to increased complementarity among species (Cardinale *et al*. 2007, Fargione *et al*. 2007, Reich *et al*. 2012, Meyer *et al*. 2016). It was suggested that increased complementarity may originate from evenly distributed resource depletion in mixtures or negative plant–soil feedbacks developing in monocultures (Fargione *et al*. 2007). Moreover, increased complementarity was also attributed to phenotypic plasticity (Ghalambour, McKay, Carroll, & Reznick 2007) or selection of genotypes that have an advantage to grow in mixtures, i.e., “mixture-type plants”. Indeed, recent common-environment experiments with plant material from a grassland biodiversity experiment (the Jena Experiment, Roscher *et al*. 2004) suggest that increased biodiversity effects have a heritable component (Zuppinger-Dingley *et al*. 2014, van Moorsel, Schmid, Hahl, Zuppinger-Dingley, & Schmid 2018c). Plants originating from mixed communities showed stronger complementarity effects than plants originating from monoculture communities if they were grown in two-species mixtures in the glasshouse, indicating that community composition can lead to phenotypic trans-generational effects (Zuppinger-Dingley *et al*. 2014, Rottstock, Kummer, Fischer, & Joshi 2017, van Moorsel *et al*. 2018b). However, it remains unclear whether the trans-generational effects observed in these studies were due to genetic differentiation, epigenetic differences or (possibly epigenetically-induced) maternal effects (Tilman & Snell-Rood 2014).

While phenotypic changes have been widely linked to genetic variation, an increasing body of evidence suggests epigenetic mechanisms to play an important role in phenotypic variation, and hence ecological processes (e.g., Bird 2007, Bossdorf, Richards, & Pigliucci 2008, Niederhuth & Schmitz 2014, Verhoeven, Vonholdt, & Sork 2016). Recent work on epigenetic recombinant inbred lines (epiRILs) of *Arabidopsis thaliana* indeed suggests a considerable contribution of induced epialleles to phenotypic variation, which is independent of genetic variation (Latzel *et al*. 2013, Cortijo *et al*. 2014, Kooke *et al*. 2015). Schmitz et al. (2013) further found evidence for epigenomic diversity which was potentially independent of genetic diversity in natural *Arabidopsis thaliana* successions. However, the importance of epigenetics in natural populations, in particular of non-model species, and whether it contributes to adaptation remains elusive (Quadrana & Colot, 2016, Richards *et al*. 2017, Groot *et al*. 2018). For example, Dubin et al. (2015) found that differences in DNA methylation between natural populations of *A. thaliana* were largely due to *trans*-acting loci, many of which showed evidence of local adaptation. Nonetheless, a recent selection experiment with *A. thaliana* suggests that epigenetic variation may indeed contribute to rapid heritable changes and adaptation (Schmid *et al*. 2018a).

Here, we tested whether community diversity can act as a selective environment resulting in genetic or epigenetic divergence. In an earlier experiment by van Moorsel *et al*. (2018b), phenotypic differences between offspring from plants that were selected in mixtures versus monocultures in the Jena Experiment were recorded when reciprocally grown in monocultures or mixtures. In the present study, we hypothesize that these phenotypic differences between plant populations within several grassland species are caused by genetic and additional epigenetic differentiation. Genetic differences were quantified as differences in DNA sequence (single nucleotide polymorphisms; SNPs). To assess epigenetic variation across plant individuals, we looked at levels of DNA cytosine methylation, which is a well-studied inheritable epigenetic modification involved in a large number of biological processes (Law & Jacobsen, 2010).

## 2. Material and Methods

### 2.1. Plant selection histories

To test whether plant communities that were grown in either monocultures or mixtures, showed genetic or epigenetic differentiation, material from plant populations from a large biodiversity field experiment (the Jena Experiment, Jena, Thuringia, Germany, 51 °N, 11 °E, 135 m a.s.l., see Roscher *et al*. 2004 and Weisser *et al*. 2017 for experimental details) were used (see also Fig. 1).

**Figure 1.**
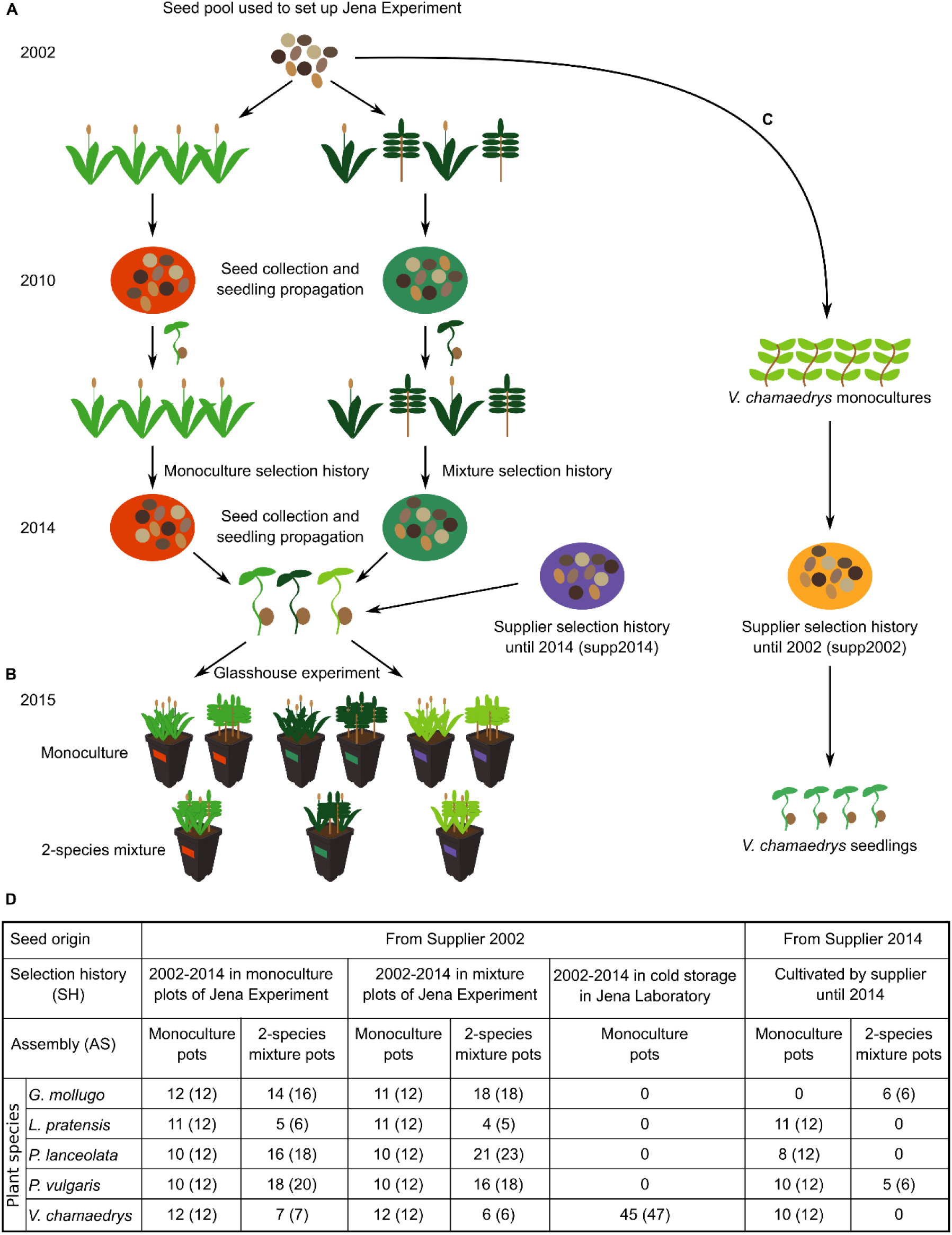
Overview of the experiment. Details are provided in the Material and methods section. (A) The origin of seeds used for the glasshouse experiment and genetic analysis. Seedlings were planted in mixtures and monocultures in the Jena Experiment in the year 2002 (Weisser *et al*. 2017). Two reproduction events occurred when seeds were collected, and subsequently new seedlings were produced and planted again in the same community composition. (B) Schematic representation of the glasshouse experiment. Monoculture assemblies and two-species mixture assemblies were planted with either plants with mixture selection history (green), monoculture selection history (orange) or supp2014 plants originating from a commercial seed supplier (blue). (C) Seeds from offspring of the original seed pool of the Jena Experiment (supp2002) were grown in an experimental garden. Figure modified after van Moorsel *et al*., 2018c. (D) Table with the experimental design. Numbers in parenthesis equal to the number of sequenced individuals. Smaller numbers in front of the parenthesis correspond to the number of individuals used during all analyses.

In the original design at Jena, 16 plant species were present in large 20 × 20 m monoculture and mixture plots from which cuttings were harvested after 8 years of growth in either mono- or mixed cultures. Out of the 16 species, four grew poorly and for several of the remaining 12, seed collection was limited (van Moorsel et al. 2018c). Hence, we were restricted to the following five species (van Moorsel et al. 2018c): the three small herbs *Plantago lanceolata* L., *Prunella vulgaris* L. and *Veronica chamaedrys* L., the tall herb *Galium mollugo* L. and the legume *Lathyrus pratensis* L for subsequent propagation and reciprocal treatments.

To gauge the differences between plants grown in the Jena Experiment and plants that experienced a different selection environment, we obtained seeds from the original seed supplier of the Jena Experiment (Rieger Hofmann GmbH, in Blaufelden-Raboldshausen, Germany) as outgroups. To test how similar these outgroup seeds were to the original seed pool that was used to set up the Jena Experiment in 2002, we also used seed material from the original seed pool. However, this was only possible for one species, *V. chamaedrys*. According to the seed supplier, all seeds were harvested from plants that were originally collected at different field sites in Germany and then propagated for at least five years in monocultures with reseeding them every year (van Moorsel *et al*. 2018a). Although this does not guarantee close similarity with the original seed pool that was used at the start of the Jena Experiment, it does provide good material to test the difference between plants that have grown in the Jena Experiment and those that experienced different selective environments.

In summary, there were three selection histories for all species and an additional fourth history for *V. chamaedrys* (see also Fig. 1): 1) monoculture in Jena, 2) mixture in Jena, 3) monoculture in the fields of the seed supplier until 2014 and 4) monoculture in the fields of the seed supplier until 2002 (only for *V. chamaedrys*). Histories 3) and 4) will be abbreviated as supp2014 and supp2002 (Tab. S1).

### 2.2. Seed collection in monoculture and mixture histories

Given that all plant species used in the study are perennial plants, it is possible that they reproduced mostly vegetative in the field. Therefore, plants with a selection history in either mixture or monoculture in the Jena Experiment underwent two controlled reproductive cycles in 2010 and 2014. This additional step aimed to increase the genetic diversity, which can then be subject to selection, and to reduce maternal effects and carry-overs of somatic epigenetic marks.

In spring 2010, cuttings from all plant communities were collected and transplanted to an experimental garden in Zurich, Switzerland, in an identical plant composition as in the Jena Experiment, for the first controlled pollination and seed production (see also Zuppinger-Dingley *et al*. 2014). In spring 2011, the seedlings produced from these seeds were transplanted back to the same plots of the Jena Experiment from where the parents had originally been collected and in the same community composition (see Tab. S2 for the community compositions of the plots in the Jena Experiment). In March 2014, plant communities of the plots that were re-established in 2011 in the Jena Experiment were again transferred to the experimental garden in Zurich for the second controlled pollination and seed production. For each experimental plot, we excavated several blocks of soil including the entire vegetation (in total one square meter). These blocks were then used to establish the plots in the experimental garden. We added a 30 cm layer of soil (1:1 mixture of garden compost and field soil, pH 7.4, commercial name Gartenhumus, RICOTER Erdaufbereitung AG, Aarberg, Switzerland) to each plot to make sure the plants established. During the controlled pollination and seed production plots were surrounded by nets and only left open on top to allow pollinator access. This did not fully exclude the possibility of cross-pollination between plots containing different plant communities, and such cross-pollination may also have occurred in the field during sexual reproduction events. However, such cross-pollination would have resulted in the populations becoming more similar to each other and hence, would have reduced the possibility to find genetic or epigenetic divergence. The experimental set up and design are schematically shown in Fig. 1.

### 2.3. Glasshouse experiment

The glasshouse experiment included three of the four selection histories described above (monoculture, mixture and supp2014) and an assembly treatment which corresponded to plants being planted in the glasshouse either in monocultures or mixtures as the common test environments. Hence, the full experimental design includes five plant species, three selection histories and two assembly treatments. The fourth history of *V. chamaedrys* (supp2002) was an extension of the experiment and plants were grown separately in a glasshouse in the Netherlands at a later time point (see section 2.4 further below).

#### 2.3.1. Setup of the glasshouse experiment

Seeds from monocultures, mixtures and the seed supplier (supp2014) were germinated in December 2014 in germination soil (“Anzuchterde”, Ökohum, Herbertingen, Germany) under constant conditions in the glasshouse without additional light. Seedlings were planted as monocultures of four individuals or two-species mixtures (2 × 2 individuals) into 2-L pots filled with agricultural soil (50 % agricultural sugar beet soil, 25 % perlite, 25 % sand; Ricoter AG, Aarberg, Switzerland). Species pairs in the mixtures were chosen according to seedling availability and single pots always contained four plants of the same selection history (i.e., there was no competition between different selection histories).

The experiment was replicated in six blocks, each including the full experimental design. Within each block, pots were placed on three different tables in the glasshouse at random without reference to selection history or assembly treatment. During the experiment the pots were not moved. The plants were initially kept at day temperatures of 17–20 °C and night temperatures of 13–17°C without supplemental light. To compensate for overheating in summer, an adiabatic cooling system (Airwatech; Bern, Switzerland) was used to keep inside temperatures constant with outside air temperatures.

#### 2.3.2. Phenotype measurements

The following traits were measured: plant height, leaf thickness, specific leaf area (SLA) and aboveground biomass. These traits were shown to relate to competitive growth and affect plant community productivity in biodiversity experiments (Roscher *et al*. 2015, Cadotte 2017). All traits were measured after twelve weeks from 18 May to 4 June 2015. Leaf thickness was measured for three representative leaves using a thickness gauge.

Specific leaf area (SLA) of up to 20 representative leaves (depending on the leaf size of the species) of each species in a pot was measured by scanning fresh leaves with a Li-3100 Area Meter (Li-cor Inc., Lincoln, Nebraska, USA) immediately after harvest and determining the mass of the same leaves after drying. All four individuals in a pot were sampled.

#### 2.3.3. Sampling of plant material

Samples for epigenetic and genetic analysis were harvested between 18 and 28 May 2015, after twelve weeks of plant growth in the glasshouse. We chose to sequence all individuals from the first three experimental blocks. All four plants were sampled in each pot. One young leaf per plant was cut from the living plant and immediately shock-frozen in liquid nitrogen. The samples were then stored at –80 °C until further analysis.

### 2.4. Offspring of the original seed pool (fourth selection history)

For the species *V. chamaedrys*, seeds from offspring of the original seed pool used to set up the Jena Experiment (supp2002) were stored since 2002 at –20 °C and germinated in the glasshouse as described above. Seedlings were then transferred to an experimental garden and seeds were collected one year later. The additional generation in the experimental garden was used to overcome potential maternal effects due to the old age of the stored seeds. The collected seeds were then stored at 5 °C, transported to Nijmegen and germinated in the glasshouse of Radboud University Nijmegen. Individual plants were grown in individual 2 × 2-cm squares in a potting tray filled with a potting soil consisting of “Lentse potgrond” (www.lentsepotgrond.nl) under natural light conditions (16/8 hrs. day/night). No cold treatment or vernalisation was applied for germination. Individual plants were harvested and quick frozen in liquid nitrogen after 5 weeks of growth.

### 2.5. Measuring genetic and epigenetic variation with epiGBS

We measured genetic and epigenetic variation using an improved version of a recently developed reference-free bisulfite method (“epiGBS”, van Gurp *et al*. 2016). A detailed description of the improvements is given in the supplementary methods. In brief, we used an improved combination of methylation-insensitive restriction enzymes to avoid the bias previously reported in van Gurp *et al*. 2016, a “wobble” adapter facilitating the computational removal of PCR duplicates and a conversion-control nucleotide that allowed for a more efficient identification of the Watson/Crick strand. The epiGBS libraries were sequenced on 4 Illumina HiSeq 2500 lanes at the facilities of Wageningen University & Research Plant Research International. Samples from different selection histories and species were distributed among lanes to prevent lane effects. An exception were the supp2002 samples from *V. chamaedrys* which were sequenced at a later time point.

### 2.6. Data processing

De-multiplexing, *de novo* reference construction, trimming, alignment, strand-specific variant calling and methylation calling were done for each species as described in van Gurp *et al*. (2016) with the pipeline provided by the authors available on https://github.com/thomasvangurp/epiGBS. The short reference sequences (up to 250 bp long) restricted the analysis of linkage disequilibrium in the study species because these had no reference genomes available. *De novo* reference sequences were annotated with DIAMOND (protein coding genes; NCBI non-redundant proteins as reference; version 0.8.22; (Buchfink, Xie, & Huson 2015)) and RepeatMasker (transposons and repeats; Embryophyta as reference species collection; version 4.0.6; (Smit, Hubley, & Green, 2013–2015)). We summarized the transposable element and repeat classes into “transposons” comprising DNA, LTR, LINE, SINE and RC transposon, and “repeats” including satellite, telomeric satellite, simple, rRNA, snRNA, unknown and unclassified repeats. The annotation was then used to classify the genetic and epigenetic variants into the different feature contexts (e.g., to identify whether a single nucleotide polymorphism was located in a gene or a transposon). A summary of the reference sequences is given in Tab. S3. The total reference sequence length in Tab. S3 ranges from 3 to 11% of the entire genome for the five test species.

### 2.7. Genetic variation

#### 2.7.1. Visualization of genetic distances with single nucleotide polymorphisms (SNPs)

Individuals with a SNP calling rate below 90 % were *a priori* removed from the analysis of genetic variation. These were three, eleven, five, nine, and five individuals of *G. mollugo*, *P. lanceolata*, *L. pratensis*, *P. vulgaris* and *V. chamaedrys*, respectively (Tab. S1). These samples were well distributed across the experimental treatment combinations, i.e., one or two for a single experimental group, except for the seed-supplier history by monoculture assembly combination of *P. lanceolata* for which four individuals were removed. For each species, we filtered the genetic-variation data for single nucleotide polymorphisms (SNPs) sequenced in all individuals with a total coverage between 5 and 200. SNPs homozygous for either the reference or the alternative allele in more than 95 % of all individuals were removed as uninformative SNPs. We removed all SNPs located in contigs with more than 1 SNP per 50 base pairs (2 %). First, to avoid that contigs with many SNPs dominate the analysis of genetic differentiation given that SNPs of a contig are linked to each other. Second, to avoid a potentially negative impact of misalignments. Considering that the reference contigs represent only a minor fraction of the entire genome, there may be many reads originating from other locations not represented with a reference contig, which are still similar enough to (wrongly) align to the reference contig. Hence, contigs with large number of SNPs may have a higher SNP calling error rate. To assess the impact of this filter, we also performed the analyses described below (section 2.7.2) with all contigs, irrespective of the SNP rate. Even though the filter frequently removed half of all contigs, the results were similar (FDRs are provided in the figures from the analysis with the filter but not discussed further). SNP allele frequencies were scaled with the function “scaleGen” from adegenet (version 2.0.1; Jombart (2008)) and genetic distances between the individuals were visualized with t-SNE (Maaten & Hinton 2008, Maaten 2014). We calculated 100 maps starting from different random seeds and selected the map with the lowest final error. Individual maps were calculated in R with the package Rtsne (version 0.13; Maaten & Hinton 2008, Maaten 2014). Parameters for the function Rtsne were pca = FALSE, theta = 0, perplexity = 10.

#### 2.7.2. Test for genetic differentiation between populations with single nucleotide polymorphisms (SNPs)

SNP data were processed and filtered as described above. The study design included the factors “current assembly” and “selection history” with two and three levels, respectively. However, this design was incomplete in all species except *P. vulgaris* (see Fig. 1D). In addition, *V. chamaedrys* had a fourth level of selection history, the supp2002 plants, which were grown separately from all others. Given these imbalances and the most interesting comparison being between monoculture and mixture selection histories, we did not use a full factorial model (selection history crossed with assembly and species) to test for genetic differentiation. Instead, we tested for each species each factor within all levels of the other factor for genetic differentiation. Taking *P. vulgaris* as an example, we tested for genetic differentiation between selection histories within monoculture and mixture assemblies (between all three histories and between monoculture and mixture types), and between assemblies within the supp2014, monoculture- and mixture-type selection histories. For each test, we extracted the corresponding individuals and tested for genetic differentiation with the G-statistic test (Goudet, Raymont, Meeûs, & Rousset 1996, function gstat.randtest implemented in the package hierfstat, version 0.04-22, Goudet & Jombart 2015). P-values were corrected for multiple testing to reflect false discovery rates (FDR) and the significance threshold was set to an FDR of 0.01. This analysis was carried out with (1) all SNPs, (2) SNPs located within genes, and (3) SNPs located within transposons. We chose to separately test SNPs in genes and transposons because we expected that selection more likely acted on genes and that selection of transposons would primarily occur due to genetic linkage to an advantageous gene. In addition, we expected that SNP calls are more reliable within genes because many transposon families tend to be highly repetitive. To estimate the extent to which the genetic variation was caused by the differentiation between populations we calculated average (i.e., across all tested SNPs) pairwise F_ST_ values with the function pairwise.fst from the package adegenet (version 2.0.1, Jombart 2008, Tab. S4). Because many SNPs had F_ST_ values close to zero, we assumed that only few SNPs with F_ST_ values clearly larger than zero were under selection. To estimate the maximal divergence between the populations, we therefore also calculated the F_ST_ of each individual SNP and extracted the 99th percentiles (we chose the 99th percentile because this is more robust to outliers than the highest value, Tab. 1, S5 and S6).

**Table 1.** 99^th^ percentile of F_ST_ values in the data set with all SNPs. AS, assembly, SH, selection history. For SNPs within genes or transposons see Tab. S5 and S6.

| Populations included | *G. mollugo* | *L. pratensis* | *P. lanceolata* | *P. vulgaris* | *V. chamaedrys* |
| --- | --- | --- | --- | --- | --- |
| SH within monoculture AS |  | 0.131 | 0.045 | 0.346 | 0.188 |
| SH within mixture AS | 0.227 |  |  | 0.398 |  |
| Monoculture vs mixture SH within in monoculture AS | 0.154 | 0.084 | 0.029 | 0.167 | 0.179 |
| Monoculture vs mixture SH within mixture AS | 0.130 | 0.174 | 0.035 | 0.113 | 0.215 |
| AS within supp2014 SH |  |  |  | 0.115 |  |
| AS within monoculture SH | 0.067 | 0.171 | 0.030 | 0.066 | 0.039 |
| AS within mixture SH | 0.062 | 0.123 | 0.034 | 0.064 | 0.082 |
| Comparison to supp2002 (only *V. chamaedrys*) |  |  |  |  |  |
| Supp2014 within monoculture AS |  |  |  |  | 0.120 |
| Monoculture SH |  |  |  |  | 0.098 |
| Monoculture SH within monoculture AS |  |  |  |  | 0.118 |
| Monoculture SH within mixture AS |  |  |  |  | 0.132 |
| Mixture SH |  |  |  |  | 0.073 |
| Mixture SH within monoculture AS |  |  |  |  | 0.098 |
| Mixture SH within mixture AS |  |  |  |  | 0.123 |

To identify individual SNPs that may be directly under selection, we tested for outliers with BayeScan (version 2.1, Foll & Gaggiotti 2008, Fischer, Foll, Excoffier & Heckel 2011). Given that there was no genetic differentiation between assemblies, we treated plants with the same selection histories but different assemblies as a single population. Hence, the tests either included two (monoculture vs. mixture) or three (monoculture, mixture and supp2014) selection histories. For *V. chamaedrys*, we also tested each of the three selection histories (monoculture, mixture and supp2014) against the original seed pool (supp2002). SNPs were identified as significant if the false discovery rate (FDR) was below 0.05 (Tab. S7).

### 2.8. Epigenetic variation

#### 2.8.1. Characterization of genome-wide DNA methylation levels

For each species, we filtered the epigenetic variation data for cytosines sequenced in at least three individuals per population (i.e., experimental treatment combination) with a total coverage between 5 and 200. This filter is different from the one applied for the SNP data because the down-stream analyses have different requirements regarding missing data (more flexible for the DNA methylation data). To provide an overview of the genome-wide DNA methylation levels of the five species or each experimental treatment combination per species, we visualized the DNA methylation levels of all cytosines averaged across all individuals with violin plots. We also visualized the average DNA methylation level within genes, transposons, repeats and unclassified reference contigs with heatmaps. Both was done either using all sequence contexts (CG, CHG, CHH) at once or separately for each sequence context.

#### 2.8.2. Identification of differentially methylated cytosines (DMCs)

DNA methylation data were processed and filtered as described above. Variation in DNA methylation at each individual cytosine was then analysed with a linear model in R with the package DSS (version 2.24.0; Y. Park & Wu (2016)), according to a design with a single factor comprising all different experimental treatment combinations as separate levels and using contrasts to compare levels of interest (similar to the approach described for RNA-Seq in Schmid 2017 and the testing procedure described in Schmid, Giraldo-Fonseca, Smetanin & Grossniklaus 2018b). Specific groups were compared with linear contrasts and *P*-values for each contrast were adjusted for multiple testing to reflect false discovery rates (FDR, Benjamini & Hochberg 1995). Taking *P. vulgaris* as an example, we compared the three selection histories across both assemblies and within each assembly to each other. Likewise, we compared the two assemblies across all selection histories and within each selection history to each other. A cytosine was defined as differentially methylated (“DMC”, see also Schmid *et al*. 2018a) if the FDR was below 0.01 for any of the contrasts.

### 2.9. Correlation between genetic and epigenetic data

#### 2.9.1. Overall correlation

To assess the correlation between genetic and epigenetic data, we calculated between-individual distances for both data sets and tested for correlation between the distances with Mantel tests. Genetic distances between two individuals were calculated as the average distance of all per-SNP differences. Per SNP, the distance was set to 0 if all alleles were identical, 1 if all alleles were different and 0.5 if one allele was different. Epigenetic distances between two samples were calculated as the average difference in DNA methylation across all cytosines. The tests were conducted in R with the package vegan (version 2.4-4, function mantel() with 9999 permutations; Oksanen *et al*. 2017). *P*-values were corrected per species for multiple testing to reflect false discovery rates (FDR).

#### 2.9.2. Linkage of genetic and epigenetic variation

To test how much of the genetic differentiation could be attributed to selection history, and, subsequently, how much of the epigenetic (methylation) variation was associated with selection history after controlling for differences in genetic structure that may have been induced by the selection histories, we modelled the average DNA methylation level of a given reference sequence in response to the sequence context (CTXT), the assembly treatment (AS), the genotype of the reference sequence (SNP), the interaction between the sequence context and the genotype (CTXT:SNP) and the selection history (SH) fitted in this order (percent methylation ∼ CTXT + AS + SNP + CTXT:SNP + SH + CTXT:SH). We then compared this result to an alternative model in which SH and SNP were switched (percent methylation ∼ CTXT + AS + SH + CTXT: SH + SNP + CTXT:SNP). Hence, whereas the second model tests for epigenetic differentiation between selection histories irrespective of the underlying genetics, the first model tests whether there was epigenetic differentiation between selection histories that could not be explained by the underlying genetics. We only used reference sequences which passed the coverage filters described above. We further only included the monoculture and mixture histories from the Jena field because only these two were fully factorially crossed with assembly in all species. Models were calculated with the functions lm() and anova() in R (version 3.5.1). Results from all reference sequences were collected and *P*-values for each term were adjusted for multiple testing to reflect false discovery rates (FDR, Benjamini & Hochberg 1995). Note that because of different distribution and testing procedure, results from this model with an average level of DNA methylation across several cytosines cannot be directly compared with the results from the model used to test for differential DNA methylation at individual cytosines. This model can detect dependency of epigenetic variation on genetic variation within our reference contigs with a maximal size of 250 bp. Most associations between DNA sequence variation and methylation loci decay at relatively short distances (i.e., after 200 bp in *A. thaliana* or 1 kb in *A. lyrata*; Hollister *et al*. 2010). This model may thus provide good proxy for close-*cis* associations (close to each other at the same location in the genome, i.e., close enough to be on the same 250 bp reference sequence). However, far-*cis* associations (for example a transposon insertion variant which is close to the place of origin of the reference sequence but not represented in the reference sequence, i.e., too far to be on the same 250 bp reference sequence) or *trans* dependencies (effects from other loci that are not linked to the place of origin of the reference sequence) cannot be detected. As a result, by using this model, we may have potentially overestimated the proportion of epigenetic variation that is unlinked to genetic variation.

### 2.10. Relation between genotype/epigenotype and phenotype

#### 2.10.1. Overall correlation

To assess whether variation in phenotypic traits could be related to variation in genetic and epigenetic data we used a multivariate ANOVA with genetic or epigenetic distances between individuals (DIST) as a dependent variable and phenotypic traits as explanatory variables with 9999 permutations (package vegan, version 2.4-4, function adonis(); Oksanen *et al*. 2017). The formula was DIST ∼ biomass + thickness + height + SLA. An in-depth analysis of the phenotypes in response to the experimental design has already been presented in van Moorsel *et al*. (2018c).

#### 2.10.2. Association of genotypes/epigenotypes with phenotypes

To test whether individual reference sequences correlated with phenotypic variation, we separately modelled the variation in the four phenotypic traits (biomass, height, leaf thickness and SLA) in response to the genotype (SNP) and the percent DNA methylation (METH) for a given sequence context with the same data previously used to test linkage of genetic and epigenetic variation (see section 2.9.2. above). Models were calculated with the function lm() and anova() in R (version 3.5.1). We tested both fitting orders with either SNP or METH fitted first. Hence, the formulas were TRAIT ∼ SNP + METH and TRAIT ∼ METH + SNP. Results from all reference sequences were collected and *P*-values for each term were adjusted for multiple testing to reflect false discovery rates (FDR, Benjamini & Hochberg 1995).

## 3. Results

### 3.1. Genetic variation

Visualization of genetic distances between the plant individuals separated them according to their selection history in three out of five species, namely *G. mollugo*, *P. vulgaris* and *V. chamaedrys* (Fig. 2). As expected, populations did not separate according to the assembly treatment, because plants were assigned randomly to the assembly treatment. Offspring of plants from the original seed pool (supp2002) of *V. chamaedrys* showed greater variability than plants of the same species derived from the original seed pool but with 11 years of monoculture or mixture history in the Jena Experiment. In addition, the supp2002 individuals were interspersed between these two histories, indicating that individuals with a selection history in the field had undergone differential evolution away from the original seed pool. The supp2014 plants differed from the other two selection histories in *V. chamaedrys* as well as in *G. mollugo* and *P. vulgaris*, confirming their status as “outgroups” at least in these three species. To see whether the separation observed in the visualization were significant, we tested for genetic divergence between the selection histories and the assemblies with the G-statistics test (Fig. 3, S1 and S2, Goudet *et al*. (1996)). We first focus on the results without the supp2002 plants.

**Figure 2.**
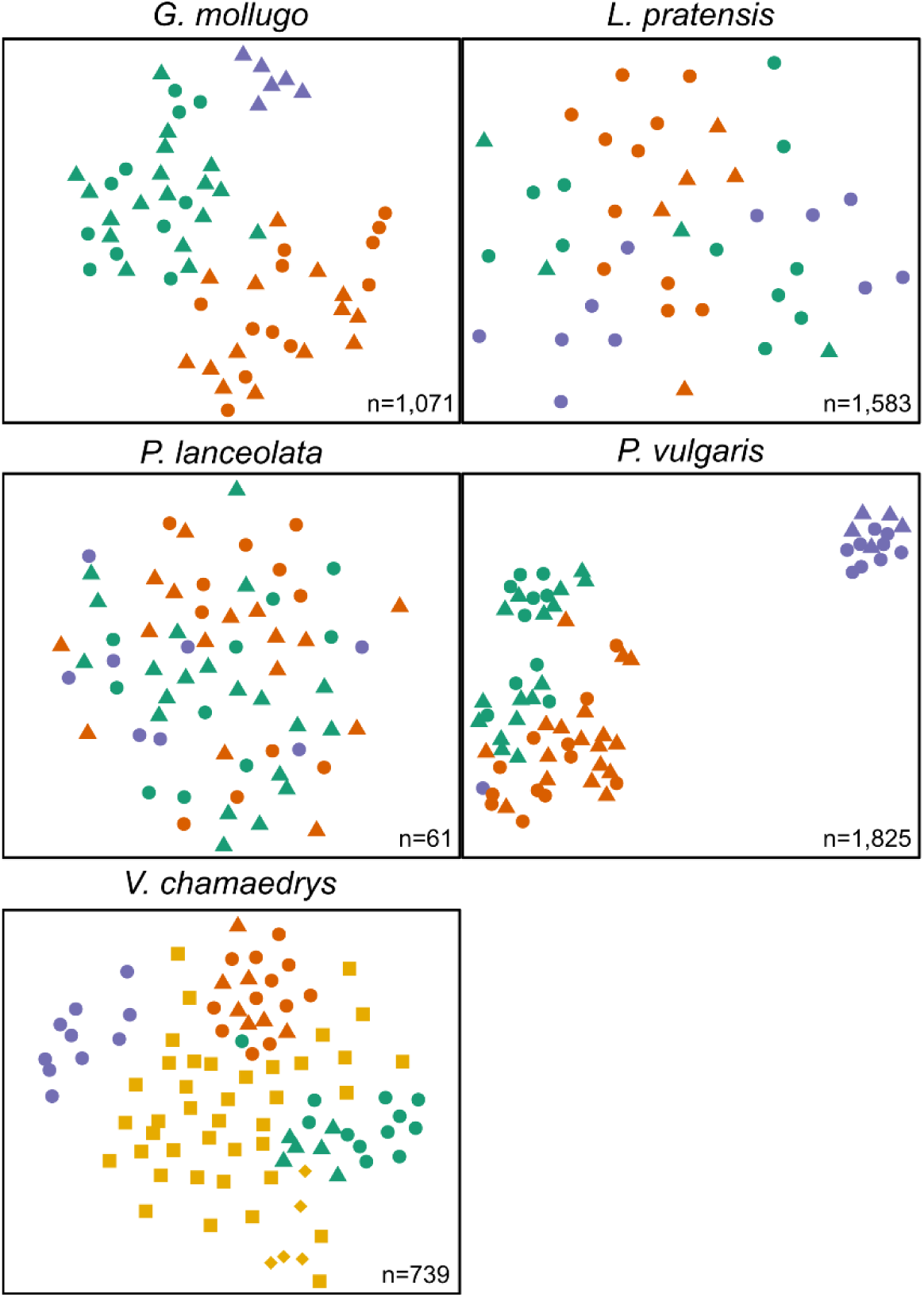
Genetic distance between individuals of the different populations for the five species. Green: selection history in mixture, orange: selection history in monocultures, blue: selection history in the field of the original seed supplier, seeds bought in 2014 (supp2014), yellow: offspring from original Jena seed pool supp2002. Triangles: monoculture assembly, circles: mixture assembly, squares: supp2002 grown in the garden, diamonds: supp2002 individuals collected from a single seed pod to qualitatively show the similarity between siblings. Assembly refers to the diversity level in the glasshouse. Note that t-SNE projection axes are arbitrary and dimensions are therefore not shown.

**Figure 3.**
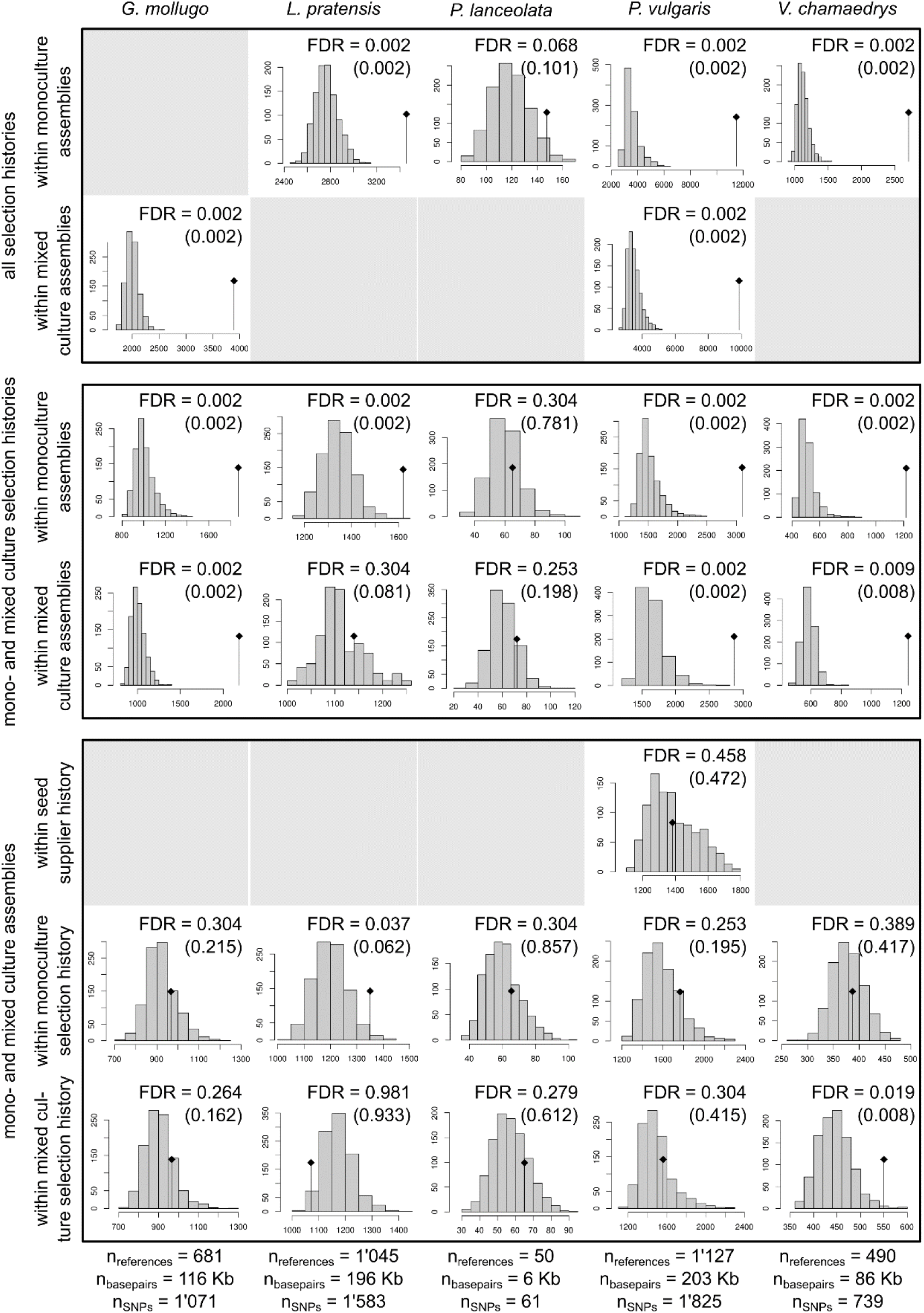
Results from the G-statistic tests given all SNPs. Each panel shows a histogram of permuted test statistics (999 permutations) and indicates the observed statistics by a black dot and a segment. Test statistics are on the x-axis, frequencies on the y-axis. Grey boxes occur where data were not available (experimental treatment combination missing). Numbers in parentheses correspond to FDRs of the same test using all reference sequences, including sequences with a SNP rate greater than 2 %.

Genetic differentiation was consistently significant (FDR < 0.01) in three of the five plant species (Fig. 3, top and middle rows). The selection histories of *P. lanceolata* did not exhibit any significant genetic differentiation. Also, the test including only the monoculture and mixture types within the mixture assemblies was not significant for *L. pratensis*. However, in *L. pratensis* statistical power was limited because there were only nine individuals available (Tab. S2, almost all other experimental groups from monoculture and mixture selection history had at least 10 individuals each). In contrast, the tests comparing the monoculture and mixture assemblies within each of the selection histories were never significant at the critical level of FDR = 0.01 (Fig. 3, bottom rows).

To estimate the amount of genetic variation explained by the selection histories, we calculated average pairwise F_ST_ values (Tab. S4) and the 99th percentiles of the SNP-wise F_ST_ values (Tab. 1, S5 and S6). Average pairwise F_ST_ values for the different selection histories were between 0.017 (supp2014 *vs.* monoculture type within the monoculture assemblies of *L. pratensis*) and 0.111 (supp2014 *vs.* mixture type within monoculture assemblies of *P. vulgaris*). With the exception of *P. lanceolata*, the 99th percentiles were markedly higher and between 0.084 (monoculture *vs.* mixture types within monoculture assemblies of *L. pratensis*) and 0.398 (all selection histories within mixture assemblies of *P. vulgaris*). Thus, overall, 1.7% to 11% of the genetic variation were explained by selection histories. However, for individual SNPs, selection histories could explain up to 40% of the genetic variation.

Within *V. chamaedrys*, comparisons between supp2002 plants and the other populations were all significant (FDR < 0.01 in all comparisons). The average pairwise F_ST_ values between the supp2002 plants and the other populations (Tab. S4) were between 0.010 and 0.015. In comparison, pairwise F_ST_ values between any of the supp2014-, monoculture-, or mixture-history populations were between 0.027 and 0.038 for this species. Likewise, the 99th percentiles of the SNP-wise F_ST_ values were consistently lower in the comparisons between the supp2002 plants and the other populations than among those (i.e., supp2014, monoculture and mixture histories populations, Tab. 1, S5 and S6). This confirmed the previous observation that supp2002 individuals, which could be considered as “parental” to the others, were genetically intermediate between the other selection histories (Fig. 2).

To identify individual SNPs that may be directly under selection, we tested for outliers with BayeScan (Tab. S7). While we could not find any outliers in *G. mollugo*, *P. lanceolata* and *V. chamaedrys*, we could identify several significant SNPs in both tests of *P. vulgaris*. 13 SNPs were significant if the three selection histories were compared with each other and 7 SNPs were significant if the monoculture and mixture selection histories were compared with each other. We could also identify a significant SNP in *L. pratensis* between the monoculture and mixture selection histories, but only if tested with all reference contigs, including the ones with a SNP rate above 2 %. These results are in parallel to the results with the 99^th^ percentiles for which *P. vulgaris* exhibited the highest F_ST_ values (Tab. 2). However, it is difficult to assess the functional relevance of these SNPs because all of them were annotated as either unknown, repeat or transposable element (data not shown).

**Table 2.** Number of cytosines with significant differences (FDR < 0.01) in DNA methylation between selection-history treatments and assemblies. AS, assembly, SH, selection history. For data on separate sequence contexts see Tab. S7 (CG), S8 (CHG), and S9 (CHH). For the results of the comparisons with the supp2002 plants (*V. chamaedrys*) see Tab. S10.

|  | *G. mollugo* | *L. pratensis* | *P. lanceolata* | *P. vulgaris* | *V. chamaedrys* | average % |
| --- | --- | --- | --- | --- | --- | --- |
| SH: mixture vs. monoculture | 5734 (0.55%) | 397 (0.07%) | 160 (0.08%) | 5240 (0.27%) | 8473 (0.8%) | 0.35% |
| % in genes | 9.57 | 1.51 | 7.5 | 9.81 | 9.21 |  |
| % in transposons | 10.85 | 31.99 | 12.5 | 10.73 | 10.39 |  |
| % in repeats | 3.82 | 5.04 | 6.25 | 6.26 | 3.76 |  |
| % in unclassified contigs | 75.76 | 61.46 | 73.75 | 73.21 | 76.64 |  |
| >> within monoculture AS | 2484 (0.24%) | 414 (0.07%) | 107 (0.06%) | 2093 (0.11%) | 4397 (0.42%) | 0.18% |
| % in genes | 8.9 | 1.93 | 10.28 | 6.93 | 7.69 |  |
| % in transposons | 11.43 | 28.99 | 11.21 | 11.71 | 9.96 |  |
| % in repeats | 3.78 | 4.59 | 5.61 | 6.64 | 4.21 |  |
| % in unclassified contigs | 75.89 | 64.49 | 72.9 | 74.73 | 78.14 |  |
| >> within mixture AS | 4039 (0.39%) | 502 (0.08%) | 1049 (0.54%) | 6797 (0.35%) | 4085 (0.39%) | 0.35% |
| % in genes | 7.65 | 2.19 | 1.91 | 8.3 | 7.81 |  |
| % in transposons | 12.01 | 29.68 | 14.59 | 14.34 | 11.8 |  |
| % in repeats | 4.43 | 5.38 | 5.24 | 6.02 | 3.89 |  |
| % in unclassified contigs | 75.91 | 62.75 | 78.27 | 71.34 | 76.5 |  |
| SH: mixture vs. supp2014 | - | - | - | 19746 (1.02%) | - | (1.02%) |
| % in genes |  |  |  | 6.36 |  |  |
| % in transposons |  |  |  | 11.14 |  |  |
| % in repeats |  |  |  | 6.63 |  |  |
| % in unclassified contigs |  |  |  | 75.87 |  |  |
| >> within monoculture AS | - | 1285 (0.21%) | 464 (0.24%) | 8352 (0.43%) | 6612 (0.63%) | 0.38% |
| % in genes |  | 1.71 | 4.53 | 6.41 | 6.79 |  |
| % in transposons |  | 31.36 | 12.28 | 10.21 | 10.41 |  |
| % in repeats |  | 4.12 | 3.45 | 6.68 | 4.05 |  |
| % in unclassified contigs |  | 62.8 | 79.74 | 76.7 | 78.75 |  |
| >> within mixture AS | 6139 (0.59%) | - | - | 13749 (0.71%) | - | 0.65% |
| % in genes | 7.27 |  |  | 6.11 |  |  |
| % in transposons | 11.86 |  |  | 11.95 |  |  |
| % in repeats | 4.71 |  |  | 6.5 |  |  |
| % in unclassified contigs | 76.17 |  |  | 75.44 |  |  |
| SH: monoculture vs. supp2014 | - | - | - | 19550<br>(1.01%) | - | (1.01%) |
| % in genes |  |  |  | 6.15 |  |  |
| % in transposons |  |  |  | 11.52 |  |  |
| % in repeats |  |  |  | 6.66 |  |  |
| % in unclassified contigs |  |  |  | 75.67 |  |  |
| >> within monoculture AS | - | 1555<br>(0.26%) | 315 (0.16%) | 6625<br>(0.34%) | 6249 (0.59%) | 0.34% |
| % in genes |  | 1.74 | 8.25 | 6.19 | 6.75 |  |
| % in transposons |  | 29.9 | 13.33 | 10.17 | 9.44 |  |
| % in repeats |  | 4.37 | 4.76 | 6.93 | 4.35 |  |
| % in unclassified contigs |  | 63.99 | 73.65 | 76.71 | 79.45 |  |
| >> within mixture AS | 4874<br>(0.47%) | - | - | 15861<br>(0.82%) | - | 0.65% |
| % in genes | 7.2 |  |  | 6.56 |  |  |
| % in transposons | 11.74 |  |  | 12.14 |  |  |
| % in repeats | 3.8 |  |  | 6.3 |  |  |
| % in unclassified contigs | 77.27 |  |  | 75 |  |  |
| AS: mixture vs. monoculture | - | - | - | 883<br>(0.05%) | - | (0.05%) |
| % in genes |  |  |  | 5.55 |  |  |
| % in transposons |  |  |  | 14.5 |  |  |
| % in repeats |  |  |  | 8.27 |  |  |
| % in unclassified contigs |  |  |  | 71.69 |  |  |
| >> within supp2014 SH | - | - | - | 1762<br>(0.09%) | - | (0.09%) |
| % in genes |  |  |  | 7.89 |  |  |
| % in transposons |  |  |  | 14.36 |  |  |
| % in repeats |  |  |  | 7.95 |  |  |
| % in unclassified contigs |  |  |  | 69.81 |  |  |
| >> within monoculture SH | 1286<br>(0.12%) | 300 (0.05%) | 255 (0.13%) | 2081<br>(0.11%) | 1308 (0.12%) | 0.11% |
| % in genes | 6.38 | 3.33 | 15.69 | 6.01 | 3.44 |  |
| % in transposons | 16.87 | 28.33 | 12.16 | 14.66 | 17.74 |  |
| % in repeats | 4.98 | 5 | 8.24 | 7.4 | 3.21 |  |
| % in unclassified contigs | 71.77 | 63.33 | 63.92 | 71.94 | 75.61 |  |
| >> within mixed culture SH | 4143<br>(0.40%) | 833 (0.14%) | 460 (0.24%) | 1522<br>(0.08%) | 1956 (0.19%) | 0.21% |
| % in genes | 6.4 | 1.08 | 3.7 | 5.39 | 5.62 |  |
| % in transposons | 13.71 | 31.21 | 10.43 | 16.1 | 13.85 |  |
| % in repeats | 5.14 | 3.84 | 6.09 | 6.44 | 3.83 |  |
| % in unclassified contigs | 74.75 | 63.87 | 79.78 | 72.08 | 76.69 |  |
| Total (percentage DMCs of tested cytosines) | 19774<br>(1.91%) | 3905<br>(0.65%) | 2223<br>(1.15%) | 45231<br>(2.34%) | 20407<br>(1.93%) | 1.60% |
| Total cytosines tested | 1034753 | 598609 | 193844 | 1929089 | 1056852 |  |

### 3.2. Epigenetic variation

To get an overview of the DNA methylation data, we visualized DNA methylation levels in percent at individual cytosines for each plant species, sequence context (CG, CHG, CHH) and genomic feature context (genes, transposons, repeats and unclassified contigs, Fig. 4). For all species, DNA methylation was generally highest in the CG context (82.6%), lower in the CHG context (59.2%), and lowest in CHH context (12.2%).

**Figure 4.**
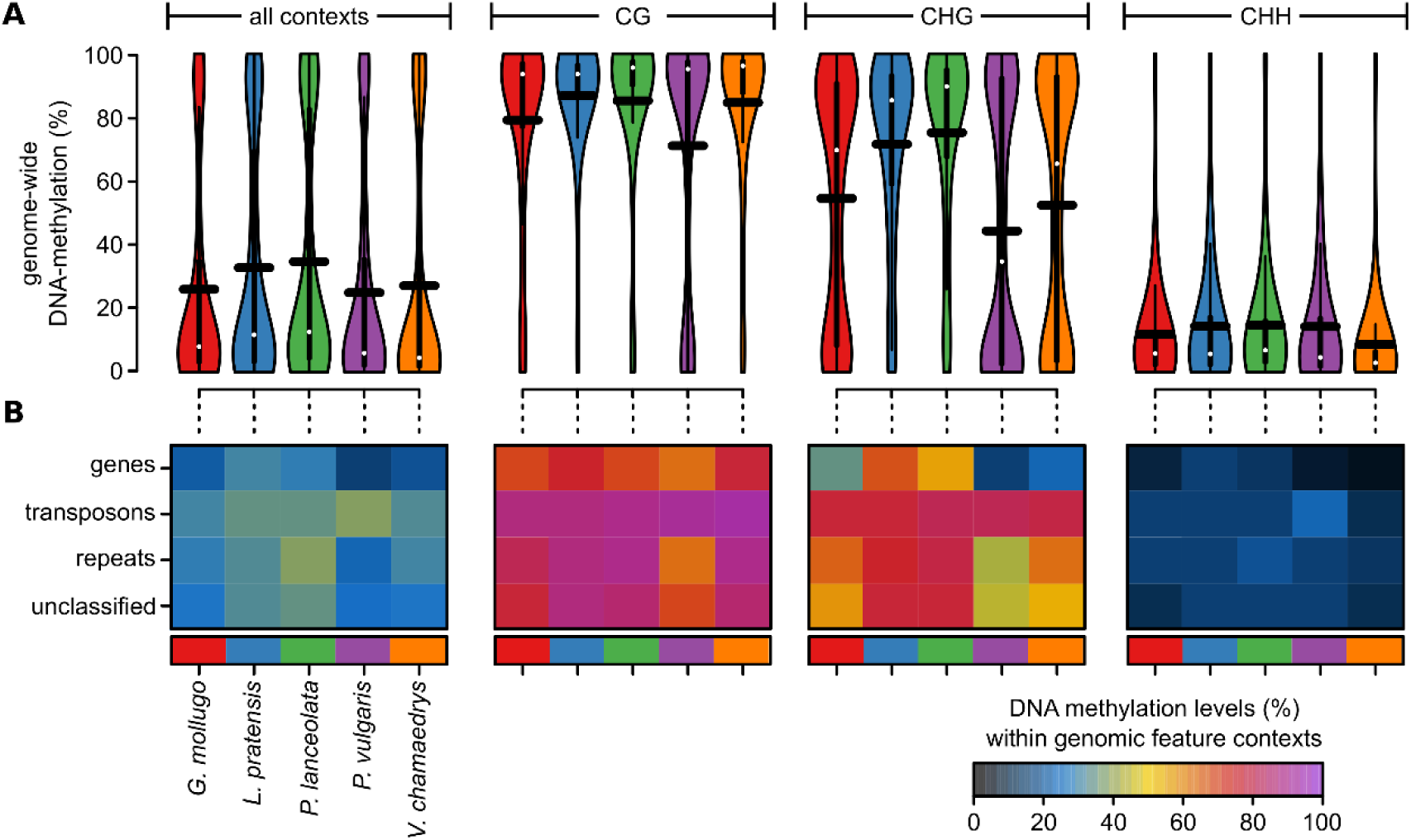
(A) DNA methylation levels in percent at individual cytosines across all or within each individual sequence context (CG, CHG, CHH) for each species used in this study shown as violin plots. The horizontal black bars correspond to the means. (B) Average DNA methylation levels in percent for each sequence context, genomic feature, and species shown as a heat map.

Differences between species were most pronounced in the CHG context in which *L. pratensis* (71.6%) and *P. lanceolata* (75.3%) exhibited markedly higher methylation levels than the other three species (54.6%, 44.4%, and 52.5% in *G. mollugo*, *P. vulgaris* and *V. chamaedrys*, respectively). Within each species and context, DNA methylation was highest in transposons and lowest in genes (Fig. 4B). Overall, these patterns are within the range of what has been reported previously for other angiosperms (e.g., Law & Jacobsen 2010, Gugger *et al*. 2016, Niederhuth *et al*. 2016, Paun, Verhoeven, & Richards 2019).

For an initial comparison between the experimental treatment combinations, we visualized the overall DNA methylation levels as we did for the different species, but for each experimental treatment combination separately (Fig. S3). Given that the overall methylation levels appeared to be highly similar between the experimental treatment combinations within species, we tested for significant differences in DNA methylation levels at each individual cytosine (Tab. 2 for all contexts and Tab. S8, S9, and S10 for each context separately). We first focused on the results excluding the supp2002 plants from *V. chamaedrys*. On average, 1.6% of all tested cytosines were significant in at least one of the tested contrasts (FDR < 0.01, “DMCs” for differentially methylated cytosine). Relative to the total number of cytosines tested, differences between selection histories (tested within or across both assemblies) were between 0.18% and 1.02% on average across all species and between 0.07% and 1.02% per individual species. Differences between the two assemblies (tested within or across all selection histories) were between 0.05% and 0.21% on average across all species and between 0.05% and 0.40% per individual species. Thus, the fraction of differentially methylated cytosines between the selection histories was generally larger than differences between the two assemblies.

Within the selection histories, differences between the monoculture types and the supp2014 plants were between 0.16% and 1.01% within species. Differences between mixture types and supp2014 plants were between 0.21% and 1.02% within species. Differences between monoculture and mixture types were between 0.06% and 0.80% within species. However, if compared within each species separately, there were always more DMCs in the comparisons between plants from Jena and the supp2014 plants than in the comparison between monoculture and mixture types. It is possible that this was at least partly due to the underlying genetic differences, given that the genetic distances between supp2014 and the other two selection histories were generally larger than the distances between the monoculture and mixture history (Tab. S4).

To further characterize the differences in DNA methylation, we calculated the average change in DNA methylation at the DMCs for each contrast, across and within all sequence contexts (CG, CHG and CHH) and feature types (genes, transposons, repeats and unclassified) and visualized these differences (Fig. 5). We could not identify clear patterns between the different comparisons with one exception: differences in the comparisons between plants from Jena and the supp2014 plants within genes (all sequence contexts) were mostly biased towards a higher methylation in the supp2014 plants. Thus, plants in the Jena Experiment showed an overall loss of DNA methylation at DMCs within genes. However, it remains unclear what functional consequences this might have had because the function of gene body methylation remains to be elucidated (Zilberman 2017).

**Figure 5.**
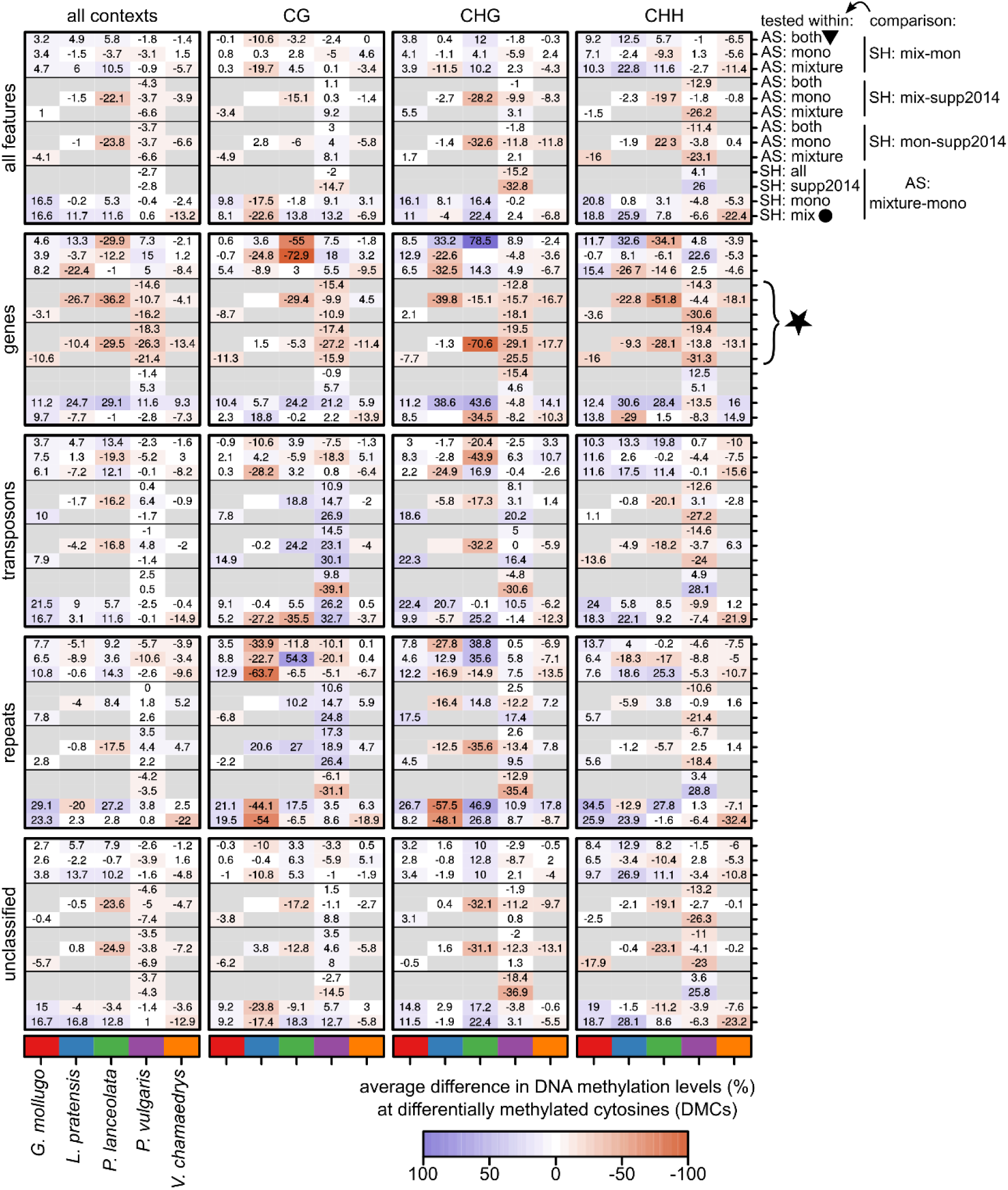
Average differences in DNA methylation at significantly differentially methylated cytosines (DMCs; FDR < 0.01) within a given sequence (all, CG, CHG, and CHH) and feature context (all, genes, transposons, repeats, unclassified) are shown for all contrasts. The comparisons and within which levels of the other factor they were tested are given in the first panel row on the right (same for all features). AS: assembly, PH: selection history. For example, the row marked with a triangle corresponds to the comparison between mixture and monoculture selection histories across both monoculture and mixture assemblies. In contrast, the row marked with a dot corresponds to the comparison between mixture assembly and monoculture assembly within the mixture selection history. The average differences are shown as colour gradient. The numbers within the heat map are the average differences. The asterisk marks the rows which show that plants in the Jena field lost on average DNA methylation at DMCs within genes compared to supp2014 plants (the two comparisons SH mix – supp2014 and SH mon – supp2014; within and across monoculture and mixture assemblies).

For *V. chamaedrys*, we also compared the supp2002 to the other experimental treatment combinations (Tab. S11). Relative to the total number of cytosines tested, differences between supp2002 plants and the other populations were between 0.82% (supp2002 *vs.* mixture history in mixture assembly) and 4.17% (supp2002 *vs.* monoculture history in both assemblies). In total, 7.4% of all cytosines tested were significant in at least one of the comparisons. Thus, even though genetically intermediate, supp2002 differed epigenetically more from the other populations than these did between each other. However, considering that these plants grew in a markedly different environment (and their ancestors had been stored as seeds for 12 years), it was not possible to separate effects of underlying genetic differences from effects of the environment. Nonetheless, it suggests that there was considerable epigenetic variation within *V. chamaedrys*, even though all plants were likely at least distantly related.

### 3.3. Correlation between genetic and epigenetic variation

To assess the correlation between genetic and epigenetic variation, we tested whether there was a significant correlation between the genetic and epigenetic distance matrices (Tab. 3). This correlation was significant (FDR < 0.05) in all species except for *G. mollugo*. Correlation to the genetic variation in these four species was highest for the CG-methylation (0.30 on average), intermediate for CHG-methylation (0.25 on average) and lowest for CHH-methylation (0.20 on average).

**Table 3.** Correlation between genetic and epigenetic variation (Pearson correlation coefficients of distance matrices). Non-significant correlations (Mantel test, FDR ≥ 0.05) are indicated by “n.s.”.

| Species | CG methylation | CHG methylation | CHH methylation |
| --- | --- | --- | --- |
| *G. mollugo* | n.s. | n.s. | n.s. |
| *L. pratensis* | 0.23 | 0.18 | 0.16 |
| *P. lanceolata* | 0.17 | 0.13 | 0.12 |
| *P. vulgaris* | 0.41 | 0.34 | 0.22 |
| *V. chamaedrys* | 0.40 | 0.36 | 0.30 |

To better estimate how much epigenetic variation was unlinked to genetic variation in close-*cis* (i.e., on the same reference sequence), we calculated the percentage of reference sequences that exhibited a significant effect of the selection history on the DNA methylation level even if an explanatory term for genotype (SNP, see section 2.9.2) was fitted first. We compared this to a model with the opposite fitting order (Tab. 4). If selection history was fitted first, its model terms SH and CTXT:SH were significant in 2.01 % of all reference sequences (average across species). However, if fitted after SNP, the effect of selection history was only significant in 0.85 % of all cases. This varied between species. For example, almost no significant effects of selection history were found in *L. pratensis* (2 out of 5,554 reference sequences) and *P. lanceolata* (1 out of 314 reference sequences) whereas up to 2.01 % of the reference sequences of *V. chamaedrys* exhibited a significant effect of selection history on DNA methylation after fitting the explanatory term for genotype first. Hence, overall and at most individual reference sequences, epigenetic variation was likely linked to genetic variation. Nonetheless, in up to 2.01 % of the reference sequences of individual species, genetic variation on the same reference sequence could not explain epigenetic variation.

**Table 4.** Percentage of reference sequences that exhibit a significant effect (FDR) in the models to test for epigenetic variation that is unlinked to genetic variation in close-*cis* (model at the bottom in which the genotype is fitted first). CTXT: sequence context of DNA methylation, AS: assembly, SH: selection history, SNP: genotype. SH & CTXT:SH and SNP & CTXT:SNP indicate the percentage of reference sequences that exhibit a significant effect in the main effect or in the interaction (union).

| Species | *G. mollugo* | *L. pratensis* | *P. lanceolata* | *P. vulgaris* | *V. chamaedrys* | average |
| --- | --- | --- | --- | --- | --- | --- |
| # Tests | 4,351 | 5,554 | 314 | 6,330 | 1,692 |  |
| Selection history fitted first |  |  |  |  |  |  |
| CTXT | 94.69 | 82.54 | 98.41 | 88.63 | 98.58 | 92.57 |
| AS | 0.39 | 0.00 | 0.32 | 0.25 | 0.00 | 0.19 |
| SH | 1.06 | 0.02 | 0.00 | 1.53 | 2.84 | 1.09 |
| CTXT:SH | 1.17 | 0.00 | 1.59 | 1.64 | 2.42 | 1.36 |
| SNP | 3.33 | 0.54 | 3.82 | 7.95 | 1.12 | 3.35 |
| CTXT:SNP | 1.40 | 1.01 | 6.69 | 5.10 | 1.30 | 3.10 |
| SNP & CTXT:SNP | 3.86 | 1.19 | 7.64 | 9.54 | 1.89 | 4.82 |
| SH & CTXT:SH | 1.86 | 0.02 | 1.59 | 2.43 | 4.14 | 2.01 |
| Genotype fitted first |  |  |  |  |  |  |
| CTXT | 95.06 | 83.72 | 98.41 | 88.67 | 98.82 | 92.94 |
| AS | 0.39 | 0.00 | 0.32 | 0.25 | 0.00 | 0.19 |
| SNP | 3.68 | 0.79 | 5.10 | 8.50 | 1.71 | 3.96 |
| CTXT:SNP | 1.79 | 1.10 | 6.37 | 5.48 | 1.65 | 3.28 |
| SH | 0.64 | 0.04 | 0.00 | 0.52 | 1.06 | 0.45 |
| CTXT:SH | 0.30 | 0.02 | 0.32 | 0.68 | 1.42 | 0.55 |
| SNP & CTXT:SNP | 4.37 | 1.42 | 7.64 | 10.19 | 2.78 | 5.28 |
| SH & CTXT:SH | 0.85 | 0.04 | 0.32 | 1.01 | 2.01 | 0.85 |

### 3.4. Relation between genetic/epigenetic variation and phenotype

To assess the relation between genetic or epigenetic variation and phenotypic variation, we tested whether phenotypic traits could explain the genetic and epigenetic distances between individuals (Tab. 5). Except for *G. mollugo*, all species had at least one phenotypic trait that could significantly (*P* < 0.05) explain genetic and sometimes also epigenetic differences. For example, leaf thickness was significant in *L. pratensis* and SLA was significant in *P. vulgaris* and *V. chamaedrys*. However, the coefficients of determination (R^2^) were with 0.02 to 0.06 relatively low, indicating that only a small fraction of the genome was correlated to the measured phenotypic traits. This was not surprising considering that we only measured few traits and that these may not have been so highly polygenic to be covered by the < 2% of the genome assessed with our reduced representation sequencing approach (i.e., epiGBS; van Gurp *et al*. 2016).

**Table 5.**
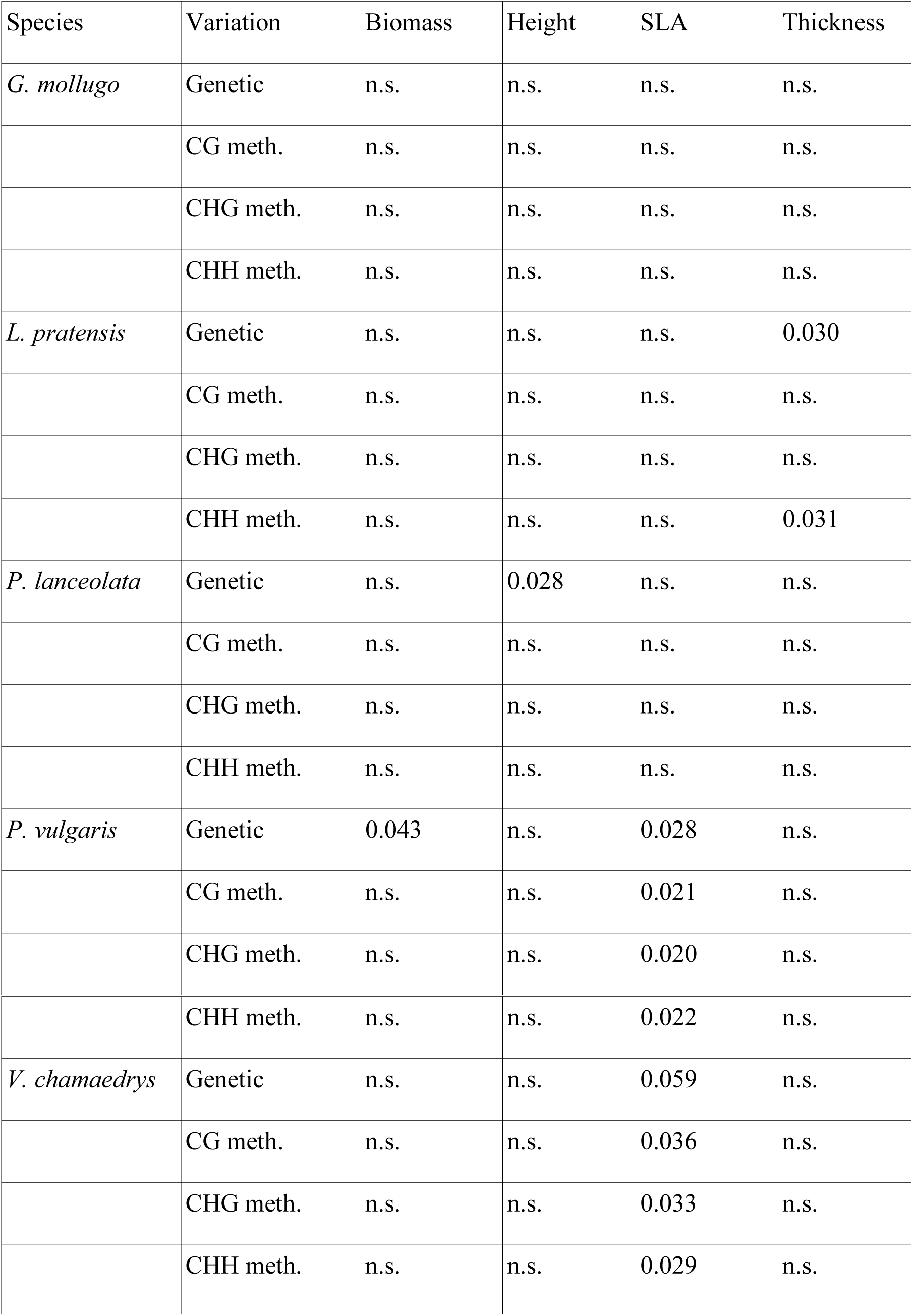
Coefficients of determination (R^2^) from multivariate ANOVAs to test whether phenotypic traits could explain genetic and epigenetic variation. Only significant (P < 0.05) results are shown. n.s.: not significant.

To identify reference sequences that were linked to the phenotypic differences, we tested for significant associations of their genotype and epigenotype with the phenotypic traits (Tab. 6). We first focused on the model in which the genotype was fitted first. All species had a trait that was at least once significantly related to genetic variation assessed with the epiGBS method (FDR < 0.05). For example, 18 and 49 reference sequences were associated with biomass in *P. vulgaris* and *V. chamaedrys*, respectively. Interestingly, *G. mollugo*, which had no significant correlations in the previous test (see Tab. 5), had a considerable amount of sequences associated with biomass or leaf thickness (429 and 320 out of 12,279, respectively). To ensure that the genetic differences in the reference sequences of *G. mollugo* were indeed also associated with the selection history, we visualized the genetic distances between the individuals (Fig. 6). The clear separation of the individuals by the factor selection history confirmed that these reference sequences were associated with the phenotype as well as the selection history.

**Figure 6.**
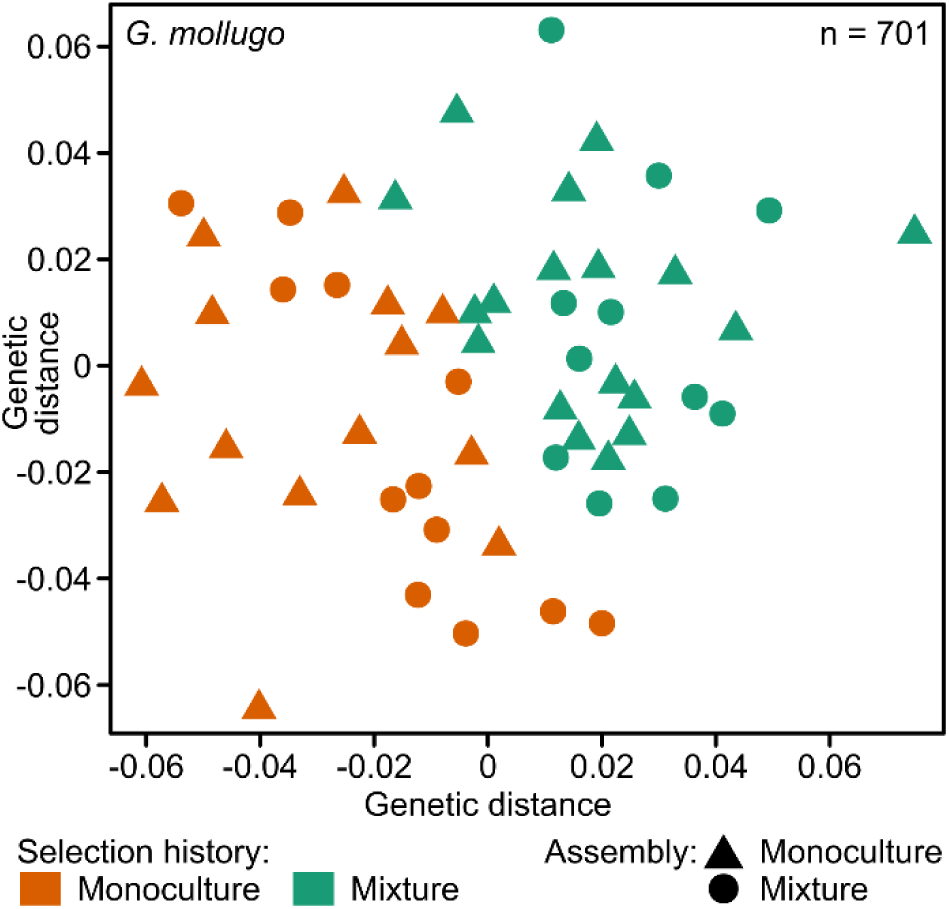
Genetic distance between the 701 reference sequences that were significantly (FDR < 0.05) associated with the phenotype in *G. mollugo*. Selection histories in this analysis were limited to the two histories in the Jena Experiment (monoculture and mixture). Distances were visualized with the function isoMDS of the R-package MASS. Genetic distances between two individuals were calculated as the average distance of all per-SNP differences. Per SNP, the distance was set to 0 if all alleles were identical, 1 if all alleles were different and 0.5 if one allele was different.

**Table 6.** Number of reference sequences with patterns of genetic (top) or epigenetic (bottom) variation that are significantly (FDR < 0.05) associated with phenotypic traits. The genotype (SNP) was fitted prior to the epigenotype (percent DNA methylation). Numbers in parenthesis correspond to the model with the inverted fitting order (TRAIT ∼ percentMethylation + SNP).

| Genotype | Biomass | Height | SLA | Thickness | # Tested |
| --- | --- | --- | --- | --- | --- |
| *G. mollugo* | 429 (79) | 0 (0) | 0 (0) | 320 (25) | 12,279 |
| *L. pratensis* | 1 (1) | 2 (1) | 0 (0) | 76 (15) | 15,797 |
| *P. lanceolata* | 0 (4) | 0 (0) | 1 (0) | 0 (0) | 904 |
| *P. vulgaris* | 18 (0) | 0 (0) | 0 (0) | 0 (0) | 17,563 |
| *V. chamaedrys* | 49 (3) | 0 (0) | 7 (1) | 0 (0) | 4,992 |
| DNA methylation | Biomass | Height | SLA | Thickness | # Tested |
| *G. mollugo* | 18 (425) | 0 (0) | 0 (0) | 16 (320) | 12,279 |
| *L. pratensis* | 0 (1) | 0 (0) | 0 (0) | 9 (73) | 15,797 |
| *P. lanceolata* | 4 (0) | 0 (0) | 0 (1) | 0 (1) | 904 |
| *P. vulgaris* | 0 (16) | 0 (0) | 0 (0) | 0 (0) | 17,563 |
| *V. chamaedrys* | 1 (41) | 0 (0) | 0 (7) | 0 (0) | 4,992 |

Epigenetic variation was rarely significantly associated with phenotypic traits if fitted after genetic variation (Tab. 6). However, if the epigenotype was fitted first, the number of reference contigs with a significant association between the epigenotype and phenotypic traits was almost identical to the number of significant associations found previously between the genotype and the phenotypic traits if the genotype was fitted first. Likewise, genetic variation was rarely significantly associated with phenotypic traits if fitted in models that already included epigenetic variation (Tab. 6). Hence, in line with the previous results, genetic and epigenetic variation that were significantly associated to phenotypic traits were both also well correlated with each other.

## 4. Discussion

For three out of five test species, namely *G. mollugo, P. vulgaris* and *V. chamaedrys*, we found genetic differences between monoculture and mixture types in a large number of SNPs. In a fourth species, *L. pratensis*, we found evidence for genetic divergence among plants grown in monoculture assemblies in the glasshouse. The comparison, however, was insignificant for plants grown in mixture assemblies, as we could only test nine individuals in total. In the fifth species, *P. lanceolata*, we could not identify significant genetic differentiation between plants with different selection histories. This finding was unexpected because *P. lanceolata* has recently been shown to exhibit clear genetic divergence after 15 years of simulated climate change (Ravenscroft, Whitlock & Fridley, 2015). It is conceivable that we could not detect genetic divergence in *P. lanceolata* because of the low number of reference sequences that passed our filter: there were only 50 sequences corresponding to 6 kb and 61 SNPs left. Thus, we might have missed regions under selection.

On average, only 1.7% to 11% of genetic variation was explained by selection histories. However, at individual SNP-level, selection histories explained up to 40% of the genetic variation. This may indicate that these loci were under selection (i.e., high divergence) whereas other parts of the genome segregated randomly (i.e, low divergence).

Besides the genetic divergence, we could also identify differences in methylation levels between the selection histories, which were generally below 1% of all tested cytosines. For *V. chamaedrys*, we observed pronounced differences in methylation levels between offspring of the original seed pool of the Jena Experiment (supp2002) and the three other selection histories. Given that these plants grew in a different environment and that their ancestors had been stored as seeds for 12 years, we could not be sure if the differences in methylation levels were due to underlying genetic or environmental differences. Nonetheless, with 7.4% of all tested cytosines being significantly differently methylated between supp2002 and the other populations (supp2014, monoculture and mixture history), there was a substantial amount of epigenetic differences within *V. chamaedrys*. Given that the supp2002 were genetically overlapping with the other groups (see Fig. 2), there was probably a considerable amount of environmentally induced epigenetic variation that was independent of genetic divergence between groups.

Overall, variation in methylation levels was significantly correlated with genetic variation in four out of five species. When we tested each reference sequence for epigenetic variation that could not be explained by genetic variation in close-*cis*, we found that up to 2.01 % of all sequences exhibited epigenetic variation that was unlinked to such genetic variation. Although this provides evidence for epigenetic divergence between selection histories that is independent and additional to genetic divergence, our analysis could not account for potential correlations between epigenetic variation and genetic variation in far-*cis* or *trans*.

We further tested to which extent the genetic and epigenetic variation was related to variation in phenotypic traits. For the genetic variation this was significant for at least one phenotypic trait in four out of five species. Epigenetic variation could also significantly explain differences in phenotypic traits in three out of four species. However, in all cases, these traits were also significantly explained by genetic variation. When we tested for associations of genetic and epigenetic variation with phenotypic traits in individual reference sequences, we could identify multiple significant associations in four out of five species. Interestingly, *G. mollugo* exhibited the highest number of associations, even though overall tests did not detect significant correlations between genetic, epigenetic or phenotypic variation. The number of significant associations between epigenetic variation and phenotypic traits were always much smaller than for genotypic variation. However, given that epigenetic variation was fitted after genetic variation, these associations suggest that some phenotypic differences might be due to epigenetic variation that is unlinked with genetic variation.

A previous selection experiment with plants found evidence for epigenetic differentiation within genotypes after few generations in *Arabidopsis thaliana* (Schmid *et al*. 2018a), but clear evidence from non-model plant species is still lacking. Our results suggest that epigenetic differences mostly reflect genetic differences and that the heritable phenotypic differences rather have a genetic than an epigenetic basis. A caveat of the novel reference-free reduced representation bisulfite sequencing method (van Gurp *et al*. 2016) is the low genome coverage (about 2 %). Thus, even if we had found more epigenetic than genetic divergence, we could not have been certain that this epigenetic divergence was unrelated to genetic divergence as we may have missed genomic regions that contain genetic loci that control for methylation. For example, in genome-wide studies with single base-pair resolution in *A. thaliana*, which revealed extensive epigenetic variation between different populations and accession, most of this variation was linked to underlying genetic differences in *cis* as well as *trans*-acting loci (Dubin *et al*. 2015, Kawakatsu *et al*. 2016, but see Schmitz et al. 2013). Such *trans*-acting loci make it difficult to separate genetics from epigenetics in non-model species because few such loci can alter large parts of the epigenome despite being only a tiny fraction of the entire genome. Hence, even though reduced representation sequencing approaches like epiGBS allow for high resolution estimates of genetic and epigenetic divergence, these techniques cannot, unfortunately, provide conclusive answers to the question whether the observed epigenetic variation has a genetic basis or not. Full exploration of the evolutionary and ecological relevance of epigenetic mechanisms may only be possible with whole-genome bisulfite sequencing and for species with high quality reference genomes (Niederhuth & Schmitz 2014; Schmid *et al*. 2018a; Paun, Verhoeven, & Richards 2019), which currently is still restricting more conclusive tests of how epigenetic variation can influence plant adaptation to natural selection.

## 5. Conclusion

Our study supports the hypothesis that the phenotypic differences observed between plant populations within several grassland species derived from the Jena Experiment, a long-term biodiversity field experiment (Zuppinger-Dingley *et al*. 2014, Hahl 2017, van Moorsel *et al*. 2018c) were caused by genetic divergence and additional epigenetic divergence. This suggests that these species can evolve rapidly in response to their biotic environment, i.e. monoculture or mixed-species communities. However, due to limitations of the novel reference-free reduced representation bisulfite sequencing method that was used to measure differences in genetic variation and levels and patterns of methylation, it was not possible to fully disentangle the genetic and epigenetic determinants of the observed rapid evolution in this grassland biodiversity experiment. Thus, despite much excitement about its potential consequences (Bossdorf *et al*. 2008, Jablonka & Raz 2009, Richards *et al*. 2010, Balao, Paun, & Alonso 2018), there is still a lack of clear evidence for the relative roles of genetic and epigenetic variation in rapid plant adaptation in nature. Further progress will be possible once more high-quality reference genomes become available to enable ecologically relevant experiments with non-model species.

## Acknowledgements

We thank T. Zwimpfer, M. Furler, D. Trujillo, D. Topalovic, E. De Luca and N. Castro for technical assistance. Keygene N.V. owns patents and patent applications protecting its Sequence Based Genotyping technologies. The University of Zurich and the University of Wageningen are licensed users. This study was supported by the Swiss National Science Foundation (grants number 147092 and 166457 to B. Schmid) and the University Research Priority Program Global Change and Biodiversity of the University of Zurich. S.J.V.M. was furthermore supported by a travel grant from the ESF Congenomics network. The Jena Experiment is funded by the German Science Foundation (DFG, FOR145, SCHM1628/5-2).

## Data accessibility

Data will be made publicly available on Zenodo (DOI 10.5281/zenodo.1167563) and SRA (accession ID SRP132258) at time of acceptance.

## Authors’ contributions

S.J.V.M, P.V. and B.S. planned and designed the study, S.J.V.M. carried out the pot experiment and collected plant material, C.A.M.W. performed the lab work and created the sequencing library and T.V.G. initially processed the sequencing data. M.W.S. processed and analysed all data and produced the figures. S.J.V.M and M.W.S. wrote the manuscript with contributions from all authors.

## Supplemental information

The supplementary information contains supplementary methods, four supplemental figures and ten supplemental tables and can be accessed online.

## References

Balao, F., Paun, O., & Alonso, C. (2018). Uncovering the contribution of epigenetics to plant phenotypic variation in Mediterranean ecosystems. Plant Biology, 20, 38–49.

Balvanera, P., Pfisterer, A. B., Buchmann, N., He, J.-S., Nakashizuka, T., Raffaelli, D., & Schmid, B. (2006). Quantifying the evidence for biodiversity effects on ecosystem functioning and services. Ecology Letters 9, 1146–1156.

Bastolla, U., Fortuna, M., Pascual-García, A., Ferrera, A., Luque, B., & Bascompte, J. (2009). The architecture of mutualistic networks minimizes competition and increases biodiversity. Nature, 458, 1018–1020.

Benjamini, Y., & Hochberg, Y. (1995). Controlling the false discovery rate: A practical and powerful approach to multiple testing. J R STAT SOC B, 57, 289–300.

Bird, A. (2007). Perceptions of epigenetics. Nature, 447, 396–398.

Bossdorf, O., Richards, C., & Pigliucci, M. (2008). Epigenetics for ecologists. Ecol Lett, 11, 106–115.

Buchfink, D., Xie, C., & Huson, D. (2015). Fast and sensitive protein alignment using DIAMOND. Nat Methods, 12, 59.60.

Cadotte, M.W. (2017). Functional traits explain ecosystem function through opposing mechanisms. Ecol Lett 29, 989–996.

Cardinale, B., Wright, J., Cadotte, M., Carroll, I., Hector, A., Srivastava, D., … Weis, J. (2007). Impacts of plant diversity on biomass production increase through time because of species complementarity. Proc Natl Acad Sci U S A, 104, 18123–18128.

Cortijo, S., Wardenaar, R., Colomé-Tatcheé, M., Gilly, A., Etcheverry, M., Labadie, K., … Johannes, F. (2014). Mapping the epigenetic basis of complex traits. Science, 343, 1145–1148.

Dubin, M. J., Zhang, P., Meng, D., Remigereau, M.-S., Osborne, E. J., Casale, F. P., … Nordborg, M. (2015). DNA methylation in *Arabidopsis* has a genetic basis and shows evidence of local adaption. eLIFE, 4, e05255.

Fargione, J., Tilman, D., Dybzinski, R., Lambers, J., Clark, C., Harpole, W., … Loreau, M. (2007). From selection to complementarity: Shifts in the causes of biodiversity-productivity relationships in a long-term biodiversity experiment. P R SOC B, 274, 871–876.

Fischer, M.C, Foll, M., Excoffier, L. & Heckel, G. (2011). Enhanced AFLP genome scans detect local adaptation in high-altitude populations of a small rodent (Microtus arvalis). Molecular Ecology, 20: 1450–1462.

Foll, M. & Gaggiotti, O.E. (2008). A genome scan method to identify selected loci appropriate for both dominant and codominant markers: A Bayesian perspective. Genetics, 180, 977–993.

Gervasi, D., & Schiestl, F. (2017). Real-time divergent evolution in plants driven by pollinators. Nat Comm, 8, 14691.

Ghalambour, C., McKay, J., Carroll, S., & Reznick, D. (2007). Adaptive versus non-adaptive phenotypic plasticity and the potential for contemporary adaptation in new environments. Funct Ecol, 21, 394–407.

Goudet, J., & Jombart, T. (2015). Hierfstat: Estimation and tests of hierarchical F-statistics. Retrieved from https://CRAN.R-project.org/package=hierfstat

Goudet, J., Raymont, M., Meeûs, T., & Rousset, F. (1996). Testing differentiation in diploid populations. Genetics, 144, 1933–1940.

Gugger, P. F., Fitz Gibbon, S., PellEgrini, M., & Sork, V. L. (2016). Species wide patterns of DNA methylation variation in Quercus lobata and their association with climate gradients. Molecular Ecology, 25, 1665–1680.

Guimarães Jr, P.R., Pires, M., Jordano, P., Bascompte, J., & Thompson, J.N. (2017). Indirect effects drive coevolution in mutualistic networks Nature, 550, 511–514.

Gurp, T. van, Wagemaker, N., Wouters, B., Vergeer, P., Ouborg, J., & Verhoeven, K. (2016). epiGBS: Reference-free reduced representation bisulfite sequencing. Nat Methods, 13, 322–234.

Groot, M.P., Wagemaker, N., Ouborg, N.J., Verhoeven, K.J.F., Vergeer, P. (2018) Epigenetic population differentiation in field- and common garden-grown *Scabiosa columbaria* plants. Ecol Evol, 8(*6*), 3505–3517.

Hahl, T. (2017). Testing for co-adaptation of plants and soil organisms in a biodiversity experiment. PhD Thesis. University of Zurich. Retrieved from Zurich Open Repository and Archive: https://doi.org/10.5167/uzh-145156

Hairston, N., Ellner, S., Geber, M., Yoshida, T., & Fox, J. (2005). Rapid evolution and the convergence of ecological and evolutionary time. Ecol Lett, 8, 1114–1127.

Hendry, A.P. (2016) Eco-evolutionary Dynamics, Princeton, New Jersey, Princeton University Press.

Hollister, J. D., Smith, L. M., Guo, Y. L., Ott, F., Weigel, D., & Gaut, B. S. (2011) Transposable elements and small RNAs contribute to gene expression divergence between Arabidopsis thaliana and Arabidopsis lyrata. Proceedings of the National Academy of Sciences, 108, 2322–2327.

Isbell, F., Craven, D., Connolly, J., Loreau, M., Schmid, B., Beierkuhnlein, C., … Eisenhauer, N. (2015). Biodiversity increases the resistance of ecosystem productivity to climate extremes. Nature, 526, 574–577.

Jablonka, E., & Raz, G. (2009). Transgenerational Epigenetic Inheritance: Prevalence, Mechanisms, and Implications for the Study of Heredity and Evolution. The Quarterly Review of Biology, 84, 131–176.

Jombart, T. (2008). Adegenet: A R package for the multivariate analysis of genetic markers. Bioinformatics, 24, 1403–1405.

Joshi, J., Schmid, B., Caldeira, M., Dimitrakopoulos, P., Good, J., Harris, R., … Lawton, J. (2001). Local adaptation enhances performance of common plant species. Ecol Lett, 4, 536–544.

Kawakatsu, T., Huang, S.-S., Jupe, F., Sasaki, E., Schmitz, R., Urich, M., … Ecker, J. (2016). Epigenomic diversity in a global collection of *Arabidopsis thaliana* accessions. Cell, 166, 492–505.

Kleynhans, E., Otto, S., Reich, P., & Vellend, M. (2016). Adaptation to elevated CO2 in different biodiversity contexts. Nat Comm, 7, 12358.

Kooke, R., Johannes, F., Wardenaar, R., Becker, F., Etcheverry, M., Colot, V., … Keurentjes, J. (2015). Epigenetic basis of morphological variation and phenotypic plasticity in *Arabidopsis thaliana*. Plant Cell, 27, 337–348.

Latzel, V., Allan, E., Bortolini Silveira, A., Colot, V., Fischer, M., & Bossdorf, O. (2013). Epigenetic diversity increases the productivity and stability of plant populations. Nat Comm, 4, 2875.

Law, J. A. & Jacobsen, S.E. (2010). Establishing, maintaining and modifying DNA methylation patterns in plants and animals. Nature Rev Gen 11(*3*), 204–220.

Lipowsky, A., Schmid, B., & Roscher, C. (2011). Selection for monoculture and mixture genotypes in a biodiversity experiment. Basic Appl Ecol, 12, 360–371.

van der Maaten, L. (2014). Accelerating t-SNE using tree-based algorithms. J Mach Lern Res, 15, 1–21.

van der Maaten, L., & Hinton, G. (2008). Visualizing data using t-SNE. J Mach Lern Res, 9, 2579–2605.

Meyer, S., Ebeling, A., Eisenhauer, N., Hertzog, L., Hillebrand, H., Milcu, A., … Weisser, W. (2016). Effects of biodiversity strengthen over time as ecosystem functioning declines at low and increases at high biodiversity. Ecosphere, 7, e01619.

van Moorsel S.J., Hahl, T., Petchey, O., Ebeling, A., Eisenhauer, N., Schmid, B., & Wagg, C. (2018a). Evolution increases ecosystem temporal stability and recovery from a flood in grassland communities. bioRxiv. Retrieved from https://dx.doi.org/10.1101/262337.

van Moorsel, S. van, Hahl, T., Wagg, C., De Deyn, G., Flynn, D., Zuppinger-Dingley, D., & Schmid, B. (2018b). Community evolution increases plant productivity at low diversity. Ecol Lett, 21, 128–137.

van Moorsel, S., Schmid, M. W., Hahl, T., Zuppinger-Dingley, D., & Schmid, B. (2018c). Selection in response to community diversity alters plant performance and functional traits. Perspect Plant Ecol Evol, 33, 51–61.

Niederhuth, C. E. & Schmitz, R.J. (2014). Inheritance of DNA methylation in plant genomes. Molecular Plant 7(*3*), 472–480.

Niederhuth, C., Bewick, A., Ji, L., Alabady, M., Kim, K., Li, Q., … Schmitz, R. (2016). Widespread natural variation of DNA methylation within angiosperms. Genome Biol, 17, 194.

Oksanen, J., Blanchet, F. G., Friendly, M., Kindt, R., Legendre, P., … McGlinn, D. (2017). Vegan: Community ecology package. Retrieved from https://CRAN.R-project.org/package=vegan.

Ouborg, N., Vergeer, P., & Mix, C. (2006). The rough edges of the conservation genetics paradigm for plants. J Ecol, 94, 1233–1248.

Paun, O., Verhoeven, K. J., & Richards, C. L. (2019). Opportunities and limitations of reduced representation bisulfite sequencing in plant ecological epigenomics. New Phytologist, 221, 738–742.

Park, Y., & Wu, H. (2016). Differential methylation analysis for BS-seq data under general experimental design. Bioinformatics, 32, 1446–1453.

Quadrana, L., & Colot, V. (2016). Plant transgenerational epigenetics. Annu Rev Genet, 50, 467–491.

Ravenscroft, C., Whitlock, R., & Fridley, J. (2015). Rapid genetic divergence in response to 15 years of simulated climate change. Glob Change Biol, 21, 4165–4176.

Reich, P., Tilman, D., Isbell, F., Mueller, K., Hobbie, S., Flynn, D., & Eisenhauer, N. (2012). Impacts of biodiversity loss escalate through time as redundancy fades. Science, 336, 589–592.

Richards, C., Alonso, C., Becker, C., Bossdorf, O., Bucher, E., Colomé-Tatché, M., … Verhoeven, K. (2017). Ecological plant epigenetics: Evidence from model and non-model species, and the way forward. Ecol Lett, 20, 1576–1590.

Roscher, C., Schumacher, J., Baade, J., Wilcke, W., Gleixner, W., Weisser, W., … Schulze, E.-D. (2004). The role of biodiversity for element cycling and trophic interactions: An experimental approach in a grassland community. Basic Appl Ecol, 5, 107–121.

Roscher, C., Schumacher, J., Schmid, B., & Schulze, E.-D. (2015). Contrasting effects of intraspecific trait variation on trait-based niches and performance of legumes in plant mixtures. PLOS ONE, 10, e01197986.

Rottstock, T., Kummer, V., Fischer, M., & Joshi, J. (2017). Rapid transgenerational effects in *Knautia arvensis* in response to plant community diversity. J Ecol, 105, 714–725.

Schmid, B. (1985). Clonal growth in grassland perennials: III. genetic variation and plasticity between and within populations of *Bellis perennis* and *Prunella vulgaris*. J Ecol, 73, 819–830.

Schmid, M. W. (2017). RNA-Seq data analysis protocol: Combining in-house and publicly available data. In A. Schmidt (Ed.), Plant germline development: Methods and protocols (pp. 309–335). New York, NY: Springer New York.

Schmid, M. W., Heichinger, C., Coman Schmid, D. Guthörl, D., Gagliardini, V., Bruggmann, R., … Grossniklaus, U. (2018a). Contribution of epigenetic variation to adaptation in *Arabidopsis*. Nat Commun, 9: 4446.

Schmid, M. W., Giraldo-Fonseca, A., Smetanin, D., & Grossniklaus, U. (2018b). Extensive epigenetic reprogramming during the life cycle of *Marchantia polymorpha*. Genome Biol, 19, 9.

Schmitz, R.J., Schultz, M.D., Urich, M.A., Nery, J.R., Pelizzola, M., Libiger, O., … Ecker, J. (2013). Patterns of population epigenomic diversity. Nature 495(*7440*), 193–198.

Schoener, T. (2011). The newest synthesis: Understanding the interplay of evolutionary and ecological dynamics. Science, 331, 426–429.

Slobodkin, L. (1961). Growth and regulation of animal populations. Holt, Rinehart; Winston, New York.

Smit, A. F. A., Hubley, R., & Green, P. (2013–2015). RepeatMasker Open-4.0. Retrieved from http://www.repeatmasker.org/

The Chimpanzee Sequencing and Analysis Consortium. (2005). Initial sequence of the chimpanzee genome and comparison with the human genome. Nature, 437, 69–87.

Tilman, D., & Snell-Rood, E. (2014). Ecology: Diversity breeds complementarity. Nature, 515, 44–45.

Verhoeven, K. J., Vonholdt, B. M., & Sork, V. L. (2016). Epigenetics in ecology and evolution: what we know and what we need to know. Molecular Ecology, 25, 1631–1638.

Weisser, W. W., Roscher, C., Meyer, S. T., Ebeling, A., Luo, G., Allan, E., … Eisenhauer, N. (2017). Biodiversity effects on ecosystem functioning in a 15-year grassland experiment: Patterns, mechanisms, and open questions. Basic and Applied Ecology, 23(*Supplement C*), 1–73.

Zilberman, D. (2017). An evolutionary case for functional gene body methylation in plants and animals. Genome Biol 18:87.

Zuppinger-Dingley, D., Schmid, B., Petermann, J., Yadav, V., De Deyn, G., & Flynn, D. (2014). Selection for niche differentiation in plant communities increases biodiversity effects. Nature, 515, 108–111.

