## Supplemental Information for "Evidence for rapid evolution in a grassland biodiversity experiment"

### Supplementary Methods

#### DNA extraction

Frozen plant material was disrupted by bead-beating frozen leaf tissue in a 2 mL eppendorf tube with 2-3 mm stainless steel beads. No more than 100 mg of fresh tissue was used per sample. DNA isolation was performed using the NucleoSpin® 8 Plant II Core Kit (740669.5 Macherey Nagel). We followed the manufacturers protocol with the following modifications. Cell lysis was achieved using Cell lysis buffer PL1 for 30 instead of 10 min. After lysis and initial centrifugation, the lysate was carefully pipetted to fresh 2.5-mL tubes, avoiding the cell debris. An extra centrifugation of 5 min at 18'000 g step was added and the lysate transferred to a 96-well rack for the next steps. To ensure efficient removal of the washing buffer after the last washing step, we centrifuged 5 min at 4'800 g and removed the remaining wash buffer. DNA concentration was measured with the Qubit® 2.0 Fluorometric dsDNA HS Assay Kit (Q32851 Life technologies).

#### DNA digestion and adapter ligation

Per individual, 30–300 ng of genomic DNA was digested overnight (17 hrs) at 37 °C in a volume of 40 µL containing 1x FD buffer (Thermo Scientific), and 2 µL of both Csp6I (FD0214, Thermo Scientific) and NsiI (R0127S, NEB). Following digestion, barcoded wobble adapters were ligated to the fragments (Figure S4). The "wobble" adapter serves to facilitate the computational removal of PCR duplicates. By adding several random nucleotides prior to the PCR, it is possible to distinguish PCR duplicates from real duplicates. As the adapter is amplified together with the sequence, PCR duplicates will have the same adapter sequence. In contrast, real duplicates have a high chance to include different wobble adapter sequences considering that there are 64 possible wobble sequences on each side of the paired-end read. To minimize the possibility of misidentifying samples as a result of sequencing or adapter synthesis error, all pair-wise combinations of barcodes differed by a minimum of three mutational steps. Barcode lengths were modulated from 4–6 bp to maximize the balance of the bases at each position in the overall set. For the ligation, 4 μL of a sample specific barcode combination of both BA and CO adapters (600 pg/μL), 6 μL T4 DNA ligase buffer, 1 μL T4 DNA ligase (M0202M, NEB) and 5 μL of distilled water were added to the digestion mix to a total volume of 60 μL. Ligation was performed for 3 hours at 22 °C followed by 4 °C overnight.

**Pooling, clean-up, and nick translation**

In order to assess the quality of libraries, the pooling was performed per species level in batches of around 12 samples per pool. When pooled, the total volume of the pool was reduced by Qiaquick PCR cleanup (28104, Qiagen) to 40 μL. The libraries were size selected by a 0.8x Agencourt AMPure XP (A63880, Beckman coulter) purification favouring > 200 bp DNA fragments and eluted in a total volume of 22 μL. To prevent the formation of adapter dimers, the barcoded adapters were not phosphorylated. Therefore, after the ligation, the insert DNA fragment –adapter connection was nicked at the 3' positions of the insert DNA fragment (Figure S4). This nick is repaired by Nick translation that recreates the total non-(5mC) methylated adapter strand and during that process also “unwobbles” the adapters since the removed nucleotides are replaced by complementing nucleotides. This nick repair prevents the partial loss of the adapter during bisulfite treatment. The nick translation reaction (1 hour at 15 °C) was performed in a reaction of 25 µL containing 19.25 µL of the purified library, 2.5 uL of 10 mM 5-methylcytosine dNTP Mix (D1030 Zymo research), 2.5 uL NEBuffer 2 and 0.75 uL DNA polymerase I (M0209, NEB).

**Bisulfite conversion**

For bisulfite conversion of non-methylated cytosines, 20 µL of the nick-translated library was used. Bisulfite treatment was performed using the EZ DNA Methylation-Lightning™ Kit (Zymo Research) with the following program according to the manufacturers protocol: 8 min at 98 °C, 1 h at 54 °C followed by up to 20 h at 4 °C.

**EpiGBS PCR**

Library amplification was performed in four individual 10-µL reactions containing 1 µL ssDNA template, 5 µL KAPA HiFi HotStart Uracil+ ReadyMix (Kapabiosystems), 3 pmol of each illumina PE PCR Primer (5’-AATGATACGGCGACCACCGAGATCTACACTCTTTCCCTACACGACGCTCTTCCGATCT-3’ and 5’-CAAGCAGAAGACGGCATACGAGATCGGTCTCGGCATTCCTGCTGAACCGCTCTTCCGATCT-3’). Temperature cycling consisted of 95 °C for 3 min followed by 18 cycles of 98 °C for 10 s, 65 °C for 15 s, 72 °C for 15 s with a final extension step at 72 °C for 5 min. Replicate PCR products were pooled and quantified using a Qubit® dsDNA HS Assay Kit (Life technologies). The quality of the Libraries was assessed by analyzing 1 µL on a High Sensitivity DNA chip on a 2100 Bioanalyzer system (Agilent). Libraries were considered suitable for sequencing if the majority of DNA fragments were between 150–400 bp. When the libraries passed quality control, they were pooled according to concentration and number of samples in the species pool, so that each individual sample was expected to yield an equal number of clusters on the Illumina flow cell. Before sequencing, the libraries were spiked with 10 % PhiX control (Illumina) to increase the complexity of the libraries.

**Sequencing**

Finally, Paired-End sequencing was performed on a Hiseq2500 sequencer using the HiSeq v4 reagents and the latest version of the HiSeq Control Software (v2.2.38), which optimizes the sequencing of low-diversity libraries (http://res.illumina.com/documents/products/technotes/technote-hiseq-low-diversity.pdf). As the first five cycles of a sequencing run are used to calculate the color matrix, our barcode design achieves almost perfect balance of the first five nucleotides when equal numbers of sequences are obtained per "A" barcode. The "B" barcodes do not have this requirement; hence same length barcodes were used. Sequencing reads were demultiplexed and deposited at SRA (SRA accession ID SRP132258). The barcode files can be accessed on Zenodo (DOI 10.5281/zenodo.1167563). In total, we generated 132'884'678, 130'721'280, 140'883'779, 215'473'088, and 194'854'086 short reads for *G. mollugo*, *L. pratensis*, *P. lanceolata*, *P. vulgaris* and *V. chamaedrys*, respectively. Read numbers for all samples and averages per treatment combination are given in Table S12 and S13.

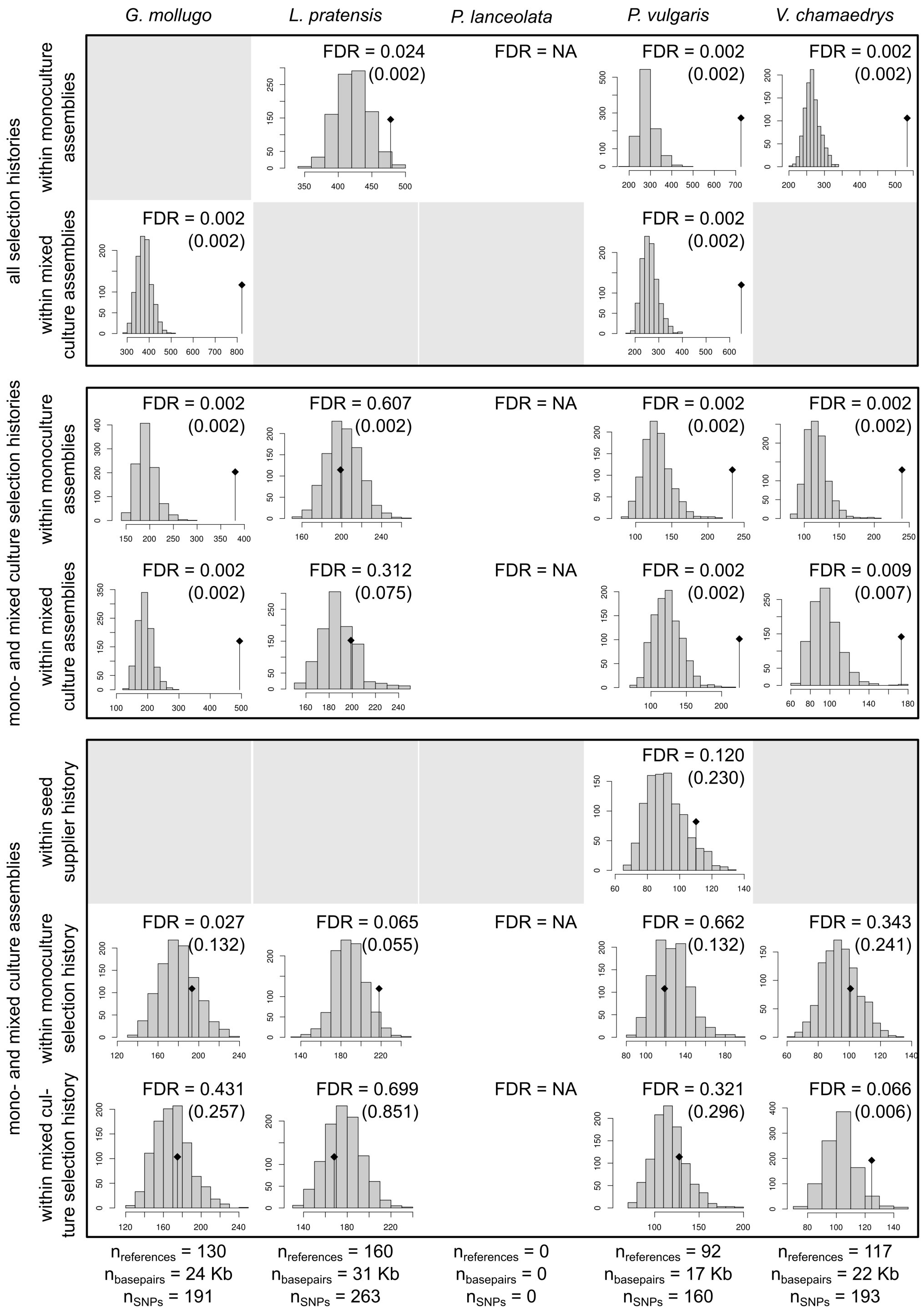

**Figure S1:** Results from the G-statistic tests given all SNPs within genes. Numbers in parentheses correspond to FDRs of the same test using all reference sequences, including sequences with a SNP rate greater than 2 %.

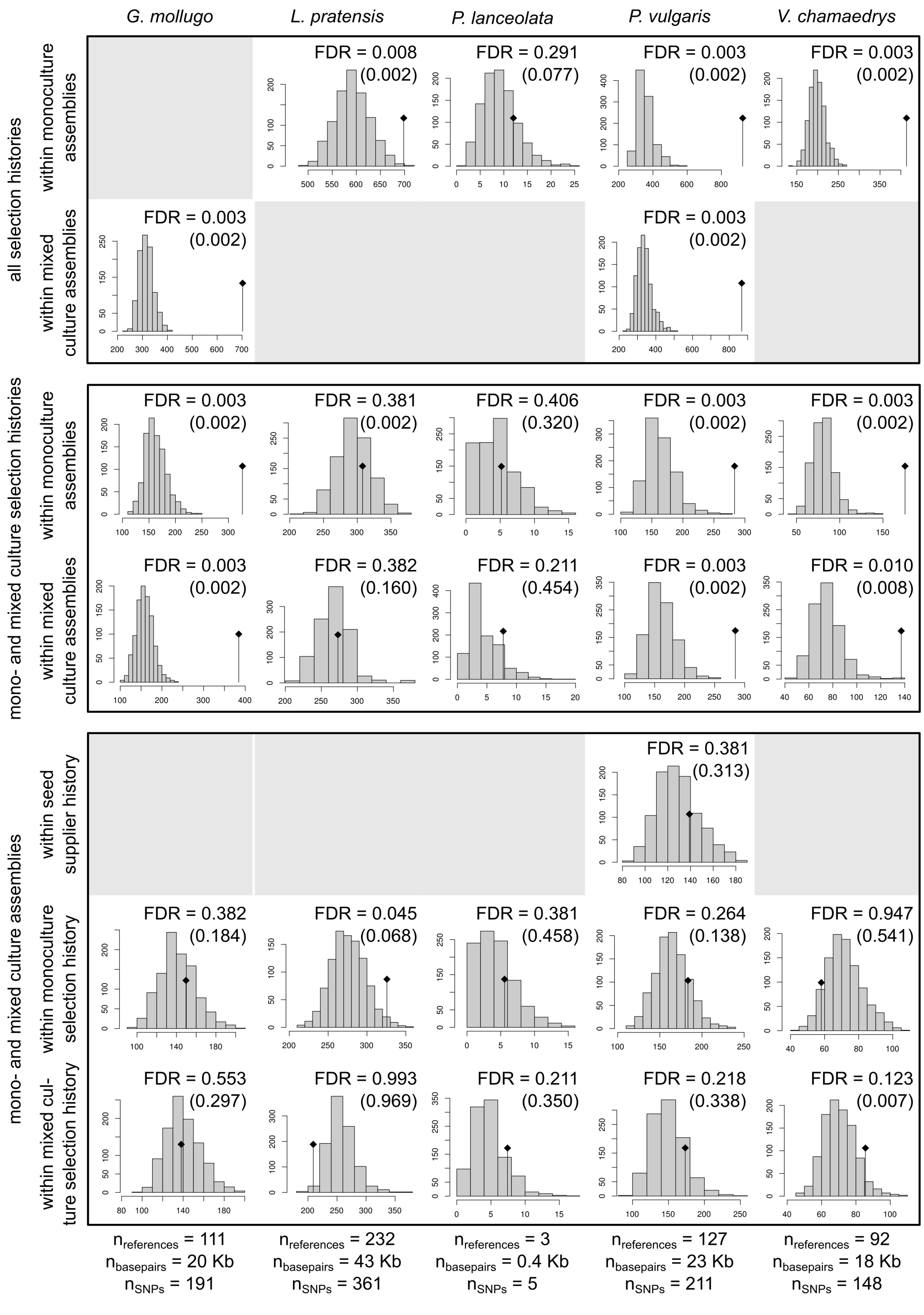

**Figure S2:** Results from the G-statistic tests given all SNPs within transposons. Numbers in parentheses correspond to FDRs of the same test using all reference sequences, including sequences with a SNP rate greater than 2 %.

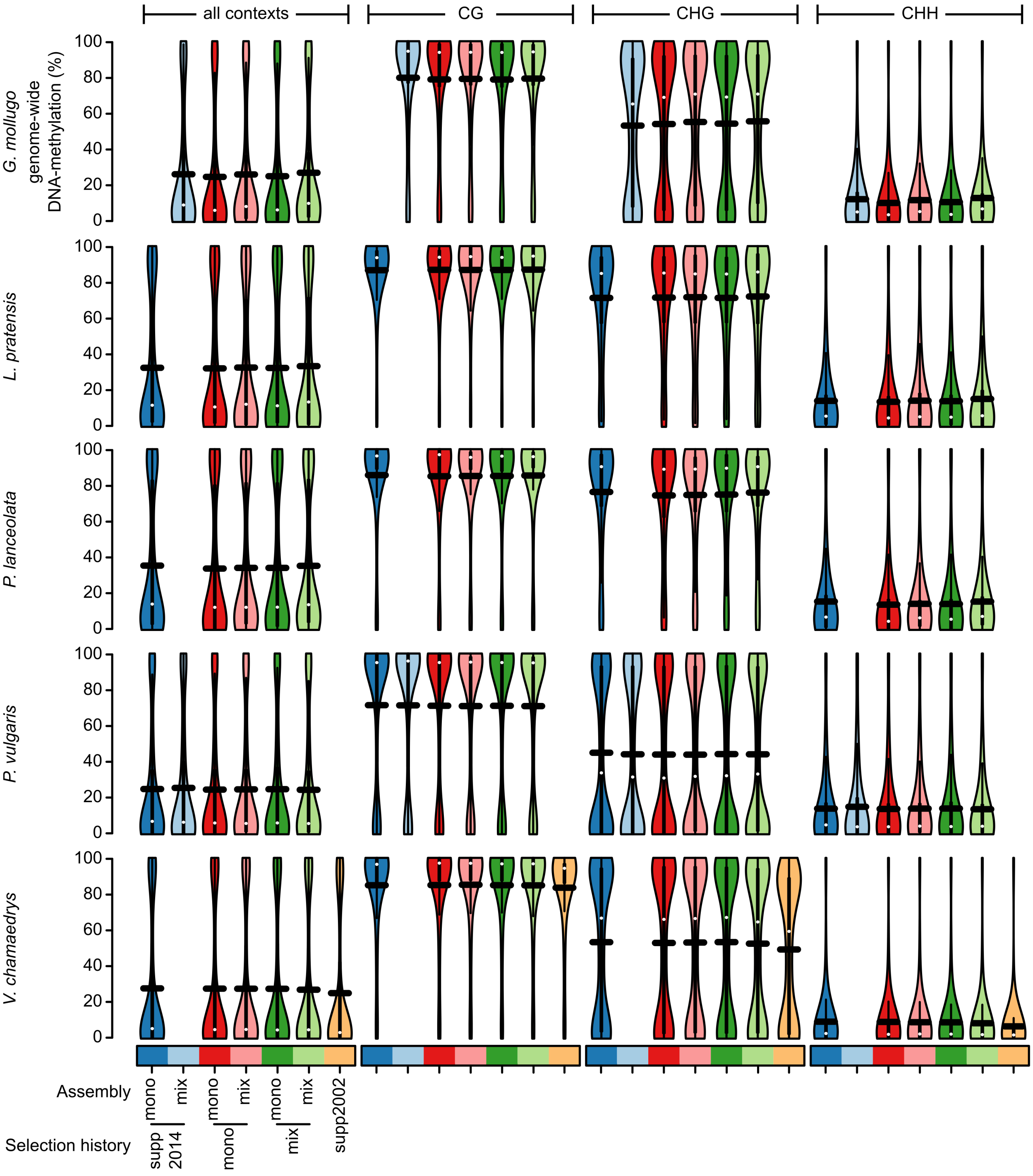

**Figure S3:** DNA methylation levels in percent at individual cytosines across all or within each individual sequence context (CG, CHG, CHH) for each experimental group of each species used.

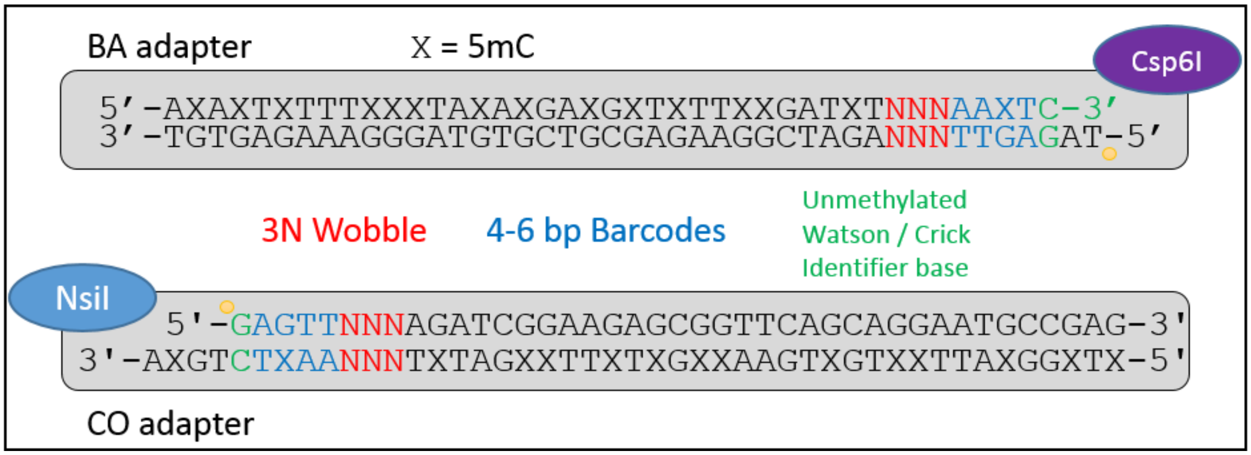

**Figure S4:** Adapter overview. Each DNA fragment – adapter connection was nicked at the position of the yellow dots.

**Table S1:** Sample overview.

| Species | Selection history | Assembly | # samples | # samples > 90% SNP calls |
|---|---|---|---|---|
| *Galium mollugo* | Monoculture | Monoculture | **12** | **12** |
| *Lathyrus pratensis* | Monoculture | Monoculture | **12** | **11** |
| *Plantago lanceolata* | Monoculture | Monoculture | **12** | **11** |
| *Prunella vulgaris* | Monoculture | Monoculture | **12** | **12** |
| *Veronica chamaedrys* | Monoculture | Monoculture | **12** | **12** |
| *Galium mollugo* | Monoculture | Mixture | **16** | **14** |
| *Lathyrus pratensis* | Monoculture | Mixture | **6** | **5** |
| *Plantago lanceolata* | Monoculture | Mixture | **18** | **16** |
| *Prunella vulgaris* | Monoculture | Mixture | **20** | **18** |
| *Veronica chamaedrys* | Monoculture | Mixture | **7** | **7** |
| *Galium mollugo* | Mixture | Monoculture | **12** | **11** |
| *Lathyrus pratensis* | Mixture | Monoculture | **12** | **11** |
| *Plantago lanceolata* | Mixture | Monoculture | **12** | **10** |
| *Prunella vulgaris* | Mixture | Monoculture | **12** | **10** |
| *Veronica chamaedrys* | Mixture | Monoculture | **12** | **12** |
| *Galium mollugo* | Mixture | Mixture | **18** | **18** |
| *Lathyrus pratensis* | Mixture | Mixture | **5** | **4** |
| *Plantago lanceolata* | Mixture | Mixture | **23** | **21** |
| *Prunella vulgaris* | Mixture | Mixture | **18** | **16** |
| *Veronica chamaedrys* | Mixture | Mixture | **6** | **6** |
| *Lathyrus pratensis* | Supp2014 | Monoculture | **12** | **11** |
| *Plantago lanceolata* | Supp2014 | Monoculture | **12** | **8** |
| *Prunella vulgaris* | Supp2014 | Monoculture | **12** | **10** |
| *Veronica chamaedrys* | Supp2014 | Monoculture | **12** | **10** |
| *Galium mollugo* | Supp2014 | Mixture | **6** | **6** |
| *Prunella vulgaris* | Supp2014 | Mixture | **6** | **5** |
| *Veronica chamaedrys* | Supp2002 | NA | **40** | **40** |
| *Veronica chamaedrys* | Supp2002 (*) | NA | **7** | **5** |

**Table S2:** Community diversity and composition of the plots the seeds originated from. Please note that each plant species grew in a different plant mixture. Hence, "plot community" refers to the mixture community the plant species grew in the Jena Experiment.

| Plot in Jena | Community diversity | # species | Target species | Plot community |
|---|---|---|---|---|
| B3A01 | Monoculture | 1 | *Galium mollugo* | *Galium mollugo* |
| B2A21 | Mix FG-Mixture | 8 | *Galium mollugo* | *Crepis biennis, Galium mollugo, Onobrychis viciifolia, Leontodon hispidus, Plantago media, Sanguisorba officinalis, Lotus cornyculatus* |
| B2A04 | Monoculture | 1 | *Geranium pratense* | *Geranium pratense* |
| B2A12 | Mono FG-Mixture | 8 | *Geranium pratense* | *Geranium pratense, Knautia arvensis, Galium mollugo, Leucanthemum vulgare, Anthriscus sylvestris, Ranunculus acris, Heracleum sphondylium, Sanguisorba officinalis* |
| B3A12 | Monoculture | 1 | *Lathyrus pratensis* | *Lathyrus pratensis* |
| B1A12 | Mono FG-Mixture | 8 | *Lathyrus pratensis* | *Lathyrus pratensis, Trifolium campestre, Trifolium dubium, Trifolium fragiferum, Trifolium hybridum, Medicago lupulina, Medicago varia, Onobrychis viciifolia* |
| B2A13 | Monoculture | 1 | *Plantago lanceolata* | *Plantago lanceolata* |
| B1A14 | Mix FG-Mixture | 8 | *Plantago lanceolata* | *Plantago lanceolata, Anthriscus sylvestris, Daucus carota, Leontodon hispidus, Luzula campestris, Trifolium campestre, Trifolium fragiferum, Trisetum flavescens* |
| B1A18 | Monoculture | 1 | *Prunella vulgaris* | *Prunella vulgaris* |
| B2A01 | Mix FG-Mixture | 4 | *Prunella vulgaris* | *Prunella vulgaris, Knautia arvensis, Trifolium pratense, Anthoxanthum odoratum* |
| B3A17 | Monoculture | 1 | *Veronica chamaedrys* | *Veronica chamaedrys* |
| B1A03 | Mix FG-Mixture | 8 | *Veronica chamaedrys* | *Veronica chamaedrys, Cyn cri, Glechoma hederacea, Lotus cornyculatus, Medicago lupulina, Phleum pratense, Primula veris, Trisetum flavescens* |

FG = functional group. The four functional groups are: legumes, tall herbs, small herbs and grasses.

**Table S3:** Reference Sequences generated in this study. Note that "genes" and "transposons/repeats" are not mutually exclusive. A reference annotated as "gene" can also be annotated as "transposon" or "repeat" (however, not both) in addition.

| annotated as | | *G. mollugo* | *L. pratensis* | *P. lanceolata* | *P. vulgaris* | *V. chamaedrys* |
|---|---|---|---|---|---|---|
| genes | number of contigs | 58'510 | 103'223 | 49'266 | 43'137 | 56'005 |
| | cummulative length (basepairs) | 11'356'223 | 19'525'195 | 9'230'439 | 8'552'287 | 10'961'834 |
| transposons | number of contigs | 67'847 | 168'470 | 37'838 | 36'897 | 48'388 |
| | cummulative length (basepairs) | 12'456'644 | 30'209'027 | 7'066'601 | 7'232'419 | 9'381'502 |
| repeats | number of contigs | 44'528 | 43'948 | 27'669 | 32'458 | 26'765 |
| | cummulative length (basepairs) | 8'328'711 | 7'974'107 | 5'303'409 | 6'430'286 | 5'182'518 |
| others | number of contigs | 672'700 | 637'396 | 442'250 | 316'399 | 543'772 |
| | cummulative length (basepairs) | 107'757'392 | 102'243'572 | 70'637'651 | 52'690'295 | 91'626'032 |
| TOTAL | number of contigs | 808'996 | 870'457 | 538'752 | 410'716 | 652'357 |
| | cummulative length (basepairs) | 133'248'303 | 144'315'524 | 88'654'861 | 71'148'267 | 112'623'385 |
| Approximative haploid genome size (basepairs) | | 1'845'000'000 1) | 4'445'000'000 1) | 1'266'880'000 2) | 635'000'000 1) | 1'329'330'000 3) |

1) **Vidiv, T, Greilhuber, J, Vilhar, B, Dermastia, M. 2009.** Selective significance of genome size in a plant community with heavy metal pollution. Ecological Applications 19: 1515-1521.

2) **Bainard, JD, Bainard, LD, Henry, TA, Fazekas, AJ, Newmaster, SG. 2012.** A multivariate analysis of variation in genome size and endoreduplication in angiosperms reveals strong phylogenetic signal and association with phenotypic traits. New Phytologist, 196: 1240-1250.

3) **Albach, DC, Geilhuber, J. 2004.** Genome Size Variation and Evolution in *Veronica*. Annals of Botany 94: 897-911

**Table S4:** Average pairwise $F_{ST}$ values. The $F_{ST}$ value measures the amount of genetic variance that can be explained by population structure (Holsinger and Weir 2009).

***G. mollugo***

| selection history | | supp2014 | | monoculture | | mixed culture | |
|---|---|---|---|---|---|---|---|
| | assembly | mono | mix | mono | mix | mono | mix |
| supp2014 | mono | | - | - | - | - | - |
| | mix | | | 0.03162 | 0.03358 | 0.03869 | 0.03017 |
| monoculture | mono | | | | 0.01207 | 0.02640 | 0.02318 |
| | mix | | | | | 0.02697 | 0.02273 |

| selection history | assembly | mono | mix | mono | mix | mono | mix |
|---|---|---|---|---|---|---|---|
| mixed culture | mono | | | | | | 0.01104 |
| | mix | | | | | | |

***L. pratensis***

| selection history | | supp2014 | | monoculture | | mixed culture | |
|---|---|---|---|---|---|---|---|
| | assembly | mono | mix | mono | mix | mono | mix |
| supp2014 | mono | | - | 0.01678 | 0.01789 | 0.01490 | 0.01865 |
| | mix | | | - | - | - | - |
| monoculture | mono | | | | 0.01812 | 0.01359 | 0.01678 |
| | mix | | | | | 0.01789 | 0.02809 |
| mixed culture | mono | | | | | | 0.01444 |
| | mix | | | | | | |

***P. lanceolata***

| selection history | | supp2014 | | monoculture | | mixed culture | |
|---|---|---|---|---|---|---|---|
| | assembly | mono | mix | mono | mix | mono | mix |
| supp2014 | mono | | - | 0.03512 | 0.01642 | 0.02220 | 0.01013 |
| | mix | | | - | - | - | - |
| monoculture | mono | | | | 0.01592 | 0.01867 | 0.01530 |
| | mix | | | | | 0.01392 | 0.00894 |
| mixed culture | mono | | | | | | 0.01057 |
| | mix | | | | | | |

***P. vulgaris***

| selection history | | supp2014 | | monoculture | | mixed culture | |
|---|---|---|---|---|---|---|---|
| | assembly | mono | mix | mono | mix | mono | mix |
| supp2014 | mono | | 0.02114 | 0.10470 | 0.09460 | 0.11099 | 0.09416 |
| | mix | | | 0.10729 | 0.08577 | 0.11395 | 0.08688 |
| monoculture | mono | | | | 0.01281 | 0.03779 | 0.02249 |
| | mix | | | | | 0.03227 | 0.01969 |
| mixed culture | mono | | | | | | 0.01237 |
| | mix | | | | | | |

***V. chamaedrys***

| Selection history | | supp2014 | | monoculture | | mixed culture | | supp2002 | |
|---|---|---|---|---|---|---|---|---|---|
| | assembly | mono | mix | mono | mix | mono | mix | garden | single seed pod * |
| supp2014 | mono | | - | 0.03368 | 0.03761 | 0.03444 | 0.03834 | 0.01306 | 0.03396 |
| | mix | | | - | - | - | - | - | - |
| monoculture | mono | | | | 0.01103 | 0.02716 | 0.03266 | 0.01452 | 0.03179 |
| | mix | | | | | 0.02848 | 0.03768 | 0.01238 | 0.04047 |
| mixed culture | mono | | | | | | 0.01656 | 0.01201 | 0.02768 |
| | mix | | | | | | | 0.00995 | 0.03937 |
| supp2002 | garden | | | | | | | | 0.00710 |
| | single seed pod | | | | | | | | |

| |
|---|
| supp2014 vs. field selection history |
| monoculture vs. mixture selection history |
| monoculture vs. mixture assemblies |
| no test for significance |

**Holsinger KE, Weir BS. 2009.** Genetics in geographically structured populations: defining, estimating and interpreting $F_{ST}$. Nature Reviews Genetics 10: 639-650.

(*) the single seed pod individuals have a relatively large $F_{ST}$ value because they are at least half-siblings and *per se* more similar to each other than random individuals of other populations. A smaller variation within a population automatically results in a larger $F_{ST}$ value because $F_{ST} = F_{total} - F_{within}$

**Table S5:** 99th percentiles of $F_{ST}$ values in the SNPs within genes. AS, assembly, SH, selection history.

| Populations included | *G. mollugo* | *L. pratensis* | *P. lanceolata (*)* | *P. vulgaris* | *V. chamaedrys* |
|---|---|---|---|---|---|
| SH within monocultury AS | | 0.1145 | | 0.31933 | 0.18695 |
| SH within mixture AS | 0.25738 | | | 0.37777 | |
| Monoculture vs mixture SH within in monoculture AS | 0.14828 | 0.0798 | | 0.1648 | 0.1892 |
| Monoculture vs mixture SH within mixture AS | 0.2134 | 0.2154 | | 0.12217 | 0.1983 |
| AS within supp2014 SH | | | | 0.10973 | |
| AS within monoculture SH | 0.082 | 0.1325 | | 0.05023 | 0.07953 |
| AS within mixture SH | 0.0569 | 0.13564 | | 0.0583 | 0.13715 |
| Comparison to supp2002 (only *V. chamaedrys*) | | | | | |
| Supp2014 within monoculture AS | | | | | 0.10634 |
| Monoculture SH | | | | | 0.1298 |
| Monoculture SH within monoculture AS | | | | | 0.12857 |
| Monoculture SH within mixture AS | | | | | 0.1719 |
| Mixture SH | | | | | 0.06683 |

| | | |
|---|---|---|
| Mixture SH within monoculture AS | | 0.07973 |
| Mixture SH within mixture AS | | 0.12842 |

(*) no genes were available with SNP info for all individuals

**Table S6:** 99th percentiles of $F_{ST}$ values in the SNPs within transposons. AS, assembly, SH, selection history.

| Populations included | *G. mollugo* | *L. pratensis* | *P. lanceolata* | *P. vulgaris* | *V. chamaedrys* |
|---|---|---|---|---|---|
| SH within monoculture AS | | 0.13086 | 0.04473 | 0.34612 | 0.18796 |
| SH within mixture AS | 0.22712 | | | 0.39801 | |
| Monoculture vs mixture SH within in monoculture AS | 0.1535 | 0.08411 | 0.02869 | 0.16692 | 0.17925 |
| Monoculture vs mixture SH within mixture AS | 0.12986 | 0.1742 | 0.03493 | 0.11295 | 0.21477 |
| AS within supp2014 SH | | | | 0.1148 | |
| AS within monoculture SH | 0.06657 | 0.17111 | 0.03005 | 0.06607 | 0.03886 |
| AS within mixture SH | 0.06198 | 0.12278 | 0.03417 | 0.06385 | 0.08207 |
| Comparison to supp2002 (only *V. chamaedrys*) | | | | | |
| Supp2014 within monoculture AS | | | | | 0.12046 |
| Monoculture SH | | | | | 0.09839 |
| Monoculture SH within monoculture AS | | | | | 0.11841 |
| Monoculture SH within mixture AS | | | | | 0.13192 |
| Mixture SH | | | | | 0.07323 |
| Mixture SH within monoculture AS | | | | | 0.09806 |
| Mixture SH within mixture AS | | | | | 0.1232 |

**Table S7:** Number of SNPs identified as outliers that are potentially under selection with BayeScan. The numbers in parentheses correspond to the number of SNPs tested. Results of the same test using all reference sequences, including sequences with a SNP rate greater than 2 % are given in the lower panel. SH, selection history. All significant SNPs were either classified as unknown, repeat or transposon (data not shown).

| Populations included | *G. mollugo* | *L. pratensis* | *P. lanceolata* | *P. vulgaris* | *V. chamaedrys* |
|---|---|---|---|---|---|
| Monoculture, mixture and supp2014 SH | 0 (1,048) | 0 (1,480) | 0 (61) | 13 (1,793) | 0 (664) |
| Monoculture vs mixture SH within mixture AS | 0 (1,040) | 0 (1,430) | 0 (61) | 7 (1,694) | 0 (608) |
| Comparison to supp2002 (only *V. chamaedrys*) | | | | | |
| Supp2014 vs supp2002 | | | | | 0 (696) |
| Monoculture vs supp2002 | | | | | 0 (697) |
| Mixture vs supp2002 | | | | | 0 (695) |
| Results with all reference sequences | | | | | |
| Monoculture, mixture and supp2014 SH | 0 (7,312) | 0 (14,830) | 0 (127) | 26 (4,240) | 0 (2,649) |
| Monoculture vs mixture SH within mixture AS | 0 (7,280) | 1 (14,488) | 0 (127) | 12 (3,988) | 0 (2,431) |
| Supp2014 vs supp2002 | | | | | 0 (2,727) |
| Monoculture vs supp2002 | | | | | 0 (2,759) |
| Mixture vs supp2002 | | | | | 0 (2,755) |

**Table S8:** Number of cytosines with significant differences (FDR < 0.01) in DNA methylation between selection history treatments and assemblies in the CG sequence context. AS: assembly, SH: selection history.

| | *G. mollugo* | *L. pratensis* | *P. lanceolata* | *P. vulgaris* | *V. chamaedrys* |
|---|---|---|---|---|---|
| SH: mixture vs. AS | 2330 | 88 | 31 | 2382 | 3969 |
| % in genes | 12.49 | 4.55 | 9.68 | 14.53 | 13.15 |
| % in transposons | 9.53 | 30.68 | 9.68 | 6.34 | 7.23 |
| % in repeats | 3.61 | 4.55 | 6.45 | 5.71 | 3.63 |
| % in unclassified contigs | 74.38 | 60.23 | 74.19 | 73.43 | 75.99 |
| >> within monoculture AS | 954 | 81 | 16 | 894 | 1932 |
| % in genes | 12.16 | 1.23 | 6.25 | 11.41 | 12.78 |
| % in transposons | 9.22 | 33.33 | 12.5 | 6.71 | 6.11 |
| % in repeats | 3.04 | 2.47 | 6.25 | 6.82 | 3.99 |
| % in unclassified contigs | 75.58 | 62.96 | 75 | 75.06 | 77.12 |
| >> within mixture AS | 1329 | 100 | 115 | 2155 | 1759 |
| % in genes | 10.46 | 3 | 3.48 | 18.79 | 12.11 |
| % in transposons | 9.1 | 36 | 12.17 | 4.64 | 7.73 |
| % in repeats | 3.84 | 5 | 4.35 | 4.5 | 4.21 |
| % in unclassified contigs | 76.6 | 56 | 80 | 72.06 | 75.95 |
| SH: mixture vs. supp2014 | - | - | - | 7099 | - |
| % in genes | | | | 10.48 | |
| % in transposons | | | | 7.96 | |
| % in repeats | | | | 6.41 | |
| % in unclassified contigs | | | | 75.15 | |
| >> within monoculture AS | - | 205 | 61 | 3325 | 2467 |
| % in genes | | 0.49 | 13.11 | 9.59 | 10.94 |
| % in transposons | | 33.17 | 6.56 | 7.28 | 7.54 |
| % in repeats | | 3.9 | 4.92 | 6.71 | 4.7 |
| % in unclassified contigs | | 62.44 | 75.41 | 76.42 | 76.81 |
| >> within mixture AS | 1963 | - | - | 4535 | - |
| % in genes | 9.17 | | | 10.39 | |
| % in transposons | 8.71 | | | 7.92 | |
| % in repeats | 4.08 | | | 6.5 | |
| % in unclassified contigs | 78.04 | | | 75.19 | |
| SH: monoculture vs. supp2014 | - | - | - | 6597 | - |
| % in genes | | | | 10.75 | |
| % in transposons | | | | 8.2 | |
| % in repeats | | | | 6.52 | |
| % in unclassified contigs | | | | 74.53 | |
| >> within monoculture AS | - | 267 | 43 | 2572 | 2496 |
| % in genes | | 0.75 | 9.3 | 9.45 | 10.94 |
| % in transposons | | 37.83 | 13.95 | 6.88 | 7.13 |

| | | | | | |
|---|---|---|---|---|---|
| % in repeats | | 3 | 2.33 | 7.62 | 4.41 |
| % in unclassified contigs | | 58.43 | 74.42 | 76.05 | 77.52 |
| >> within mixture AS | 1610 | - | - | 5062 | - |
| % in genes | 8.7 | | | 11.28 | |
| % in transposons | 7.7 | | | 7.61 | |
| % in repeats | 3.29 | | | 6.36 | |
| % in unclassified contigs | 80.31 | | | 74.75 | |
| AS: mixture vs. monoculture | - | - | - | 303 | - |
| % in genes | | | | 9.9 | |
| % in transposons | | | | 5.94 | |
| % in repeats | | | | 8.58 | |
| % in unclassified contigs | | | | 75.58 | |
| >> within supp2014 SH | - | - | - | 580 | - |
| % in genes | | | | 14.66 | |
| % in transposons | | | | 8.45 | |
| % in repeats | | | | 6.55 | |
| % in unclassified contigs | | | | 70.34 | |
| >> within monoculture SH | 317 | 48 | 35 | 532 | 296 |
| % in genes | 10.41 | 6.25 | 11.43 | 16.35 | 9.8 |
| % in transposons | 9.78 | 22.92 | 8.57 | 4.89 | 10.14 |
| % in repeats | 3.79 | 4.17 | 8.57 | 6.58 | 2.03 |
| % in unclassified contigs | 76.03 | 66.67 | 71.43 | 72.18 | 78.04 |
| >> within mixed culture SH | 408 | 132 | 51 | 435 | 531 |
| % in genes | 17.16 | 3.03 | 7.84 | 9.89 | 10.36 |
| % in transposons | 8.09 | 43.94 | 1.96 | 9.2 | 10.92 |
| % in repeats | 4.41 | 5.3 | 7.84 | 6.9 | 3.58 |
| % in unclassified contigs | 70.34 | 47.73 | 82.35 | 74.02 | 75.14 |
| Total tested | 107365 | 82338 | 23897 | 225819 | 159223 |

**Table S9:** Number of cytosines with significant differences (FDR < 0.01) in DNA methylation between selection history treatments and assemblies in the CHG sequence context. AS: assembly, SH: selection history.

| | *G. mollugo* | *L. pratensis* | *P. lanceolata* | *P. vulgaris* | *V. chamaedrys* |
|---|---|---|---|---|---|
| SH: mixture vs. monoculture | 2351 | 81 | 47 | 1257 | 2769 |
| % in genes | 8.59 | 1.23 | 2.13 | 7 | 7.19 |
| % in transposons | 9.36 | 37.04 | 4.26 | 11.22 | 13.18 |
| % in repeats | 3.28 | 1.23 | 6.38 | 6.13 | 3.32 |
| % in unclassified contigs | 78.77 | 60.49 | 87.23 | 75.66 | 76.31 |
| >> within monoculture AS | 1049 | 105 | 30 | 513 | 1409 |
| % in genes | 7.15 | 1.9 | 0 | 4.48 | 4.9 |
| % in transposons | 9.34 | 34.29 | 16.67 | 11.7 | 12.14 |
| % in repeats | 3.15 | 5.71 | 3.33 | 5.85 | 3.9 |
| % in unclassified contigs | 80.36 | 58.1 | 80 | 77.97 | 79.06 |
| >> within mixture AS | 1463 | 123 | 259 | 1242 | 1321 |
| % in genes | 6.49 | 0.81 | 2.7 | 5.96 | 5.83 |
| % in transposons | 9.57 | 39.02 | 15.83 | 15.22 | 13.47 |
| % in repeats | 4.51 | 2.44 | 4.25 | 6.84 | 3.41 |
| % in unclassified contigs | 79.43 | 57.72 | 77.22 | 71.98 | 77.29 |
| SH: mixture vs. supp2014 | - | - | - | 5879 | - |
| % in genes | | | | 5.36 | |
| % in transposons | | | | 12.59 | |
| % in repeats | | | | 5.58 | |
| % in unclassified contigs | | | | 76.48 | |
| >> within monoculture AS | - | 294 | 167 | 2838 | 2489 |
| % in genes | | 1.7 | 2.4 | 5.07 | 5.58 |
| % in transposons | | 34.01 | 13.77 | 11.06 | 10.81 |
| % in repeats | | 1.7 | 2.99 | 6.2 | 3.33 |
| % in unclassified contigs | | 62.59 | 80.84 | 77.66 | 80.27 |
| >> within mixture AS | 2734 | - | - | 3695 | - |
| % in genes | 7.39 | | | 6.06 | |
| % in transposons | 11.08 | | | 12.8 | |
| % in repeats | 4.94 | | | 5.47 | |
| % in unclassified contigs | 76.59 | | | 75.67 | |
| SH: monoculture vs. supp2014 | - | - | - | 5738 | - |
| % in genes | | | | 5.25 | |
| % in transposons | | | | 13.28 | |
| % in repeats | | | | 5.44 | |
| % in unclassified contigs | | | | 76.04 | |
| >> within monoculture AS | - | 411 | 116 | 2391 | 2322 |
| % in genes | | 2.68 | 2.59 | 5.06 | 5.3 |

| | | | | | |
|---|---|---|---|---|---|
| % in transposons | | 33.09 | 12.07 | 11.17 | 10.94 |
| % in repeats | | 3.16 | 6.03 | 6.11 | 3.53 |
| % in unclassified contigs | | 61.07 | 79.31 | 77.67 | 80.23 |
| >> within mixture AS | 2261 | - | - | 4117 | - |
| % in genes | 6.9 | | | 6.53 | |
| % in transposons | 10.79 | | | 13.63 | |
| % in repeats | 3.98 | | | 5.37 | |
| % in unclassified contigs | 78.33 | | | 74.47 | |
| AS: mixture vs. monoculture | - | - | - | 215 | - |
| % in genes | | | | 4.65 | |
| % in transposons | | | | 19.07 | |
| % in repeats | | | | 8.84 | |
| % in unclassified contigs | | | | 67.44 | |
| >> within supp2014 SH | - | - | - | 461 | - |
| % in genes | | | | 7.59 | |
| % in transposons | | | | 13.67 | |
| % in repeats | | | | 7.59 | |
| % in unclassified contigs | | | | 71.15 | |
| >> within monoculture SH | 423 | 81 | 56 | 387 | 280 |
| % in genes | 6.62 | 2.47 | 5.36 | 3.62 | 1.79 |
| % in transposons | 12.53 | 28.4 | 16.07 | 16.8 | 16.79 |
| % in repeats | 5.44 | 1.23 | 1.79 | 8.53 | 4.64 |
| % in unclassified contigs | 75.41 | 67.9 | 76.79 | 71.06 | 76.79 |
| >> within mixed culture SH | 617 | 182 | 99 | 262 | 625 |
| % in genes | 8.59 | 0 | 1.01 | 4.96 | 5.44 |
| % in transposons | 8.75 | 41.21 | 10.1 | 11.83 | 13.92 |
| % in repeats | 4.05 | 1.1 | 5.05 | 6.49 | 3.52 |
| % in unclassified contigs | 78.61 | 57.69 | 83.84 | 76.72 | 77.12 |
| Total tested | 173549 | 87668 | 35927 | 257754 | 169470 |

**Table S10:** Number of cytosines with significant differences (FDR < 0.01) in DNA methylation between selection-history treatments and assemblies in the CHH sequence context. AS: assembly, SH: selection history.

| | *G. mollugo* | *L. pratensis* | *P. lanceolata* | *P. vulgaris* | *V. chamaedrys* |
|---|---|---|---|---|---|
| SH: mixture vs. monoculture | 1053 | 228 | 82 | 1601 | 1735 |
| % in genes | 5.32 | 0.44 | 9.76 | 5 | 3.4 |
| % in transposons | 17.09 | 30.7 | 18.29 | 16.86 | 13.14 |
| % in repeats | 5.51 | 6.58 | 6.1 | 7.18 | 4.78 |
| % in unclassified contigs | 72.08 | 62.28 | 65.85 | 70.96 | 78.67 |
| >> within monoculture AS | 481 | 228 | 61 | 686 | 1056 |
| % in genes | 6.24 | 2.19 | 16.39 | 2.92 | 2.08 |
| % in transposons | 20.37 | 25 | 8.2 | 18.22 | 14.11 |
| % in repeats | 6.65 | 4.82 | 6.56 | 7 | 5.02 |
| % in unclassified contigs | 66.74 | 67.98 | 68.85 | 71.87 | 78.79 |
| >> within mixture AS | 1247 | 279 | 675 | 3400 | 1005 |
| % in genes | 6.01 | 2.51 | 1.33 | 2.5 | 2.89 |
| % in transposons | 17.96 | 23.3 | 14.52 | 20.18 | 16.72 |
| % in repeats | 4.97 | 6.81 | 5.78 | 6.68 | 3.98 |
| % in unclassified contigs | 71.05 | 67.38 | 78.37 | 70.65 | 76.42 |
| SH: mixture vs. supp2014 | - | - | - | 6768 | - |
| % in genes | | | | 2.91 | |
| % in transposons | | | | 13.22 | |
| % in repeats | | | | 7.77 | |
| % in unclassified contigs | | | | 76.09 | |
| >> within monoculture AS | - | 786 | 236 | 2189 | 1656 |
| % in genes | | 2.04 | 3.81 | 3.29 | 2.42 |
| % in transposons | | 29.9 | 12.71 | 13.57 | 14.07 |
| % in repeats | | 5.09 | 3.39 | 7.26 | 4.17 |
| % in unclassified contigs | | 62.98 | 80.08 | 75.88 | 79.35 |
| >> within mixture AS | 1442 | - | - | 5519 | - |
| % in genes | 4.44 | | | 2.63 | |
| % in transposons | 17.61 | | | 14.69 | |
| % in repeats | 5.13 | | | 7.19 | |
| % in unclassified contigs | 72.82 | | | 75.48 | |
| SH: monoculture vs. supp2014 | - | - | - | 7215 | - |
| % in genes | | | | 2.66 | |
| % in transposons | | | | 13.17 | |
| % in repeats | | | | 7.76 | |
| % in unclassified contigs | | | | 76.41 | |
| >> within monoculture AS | - | 877 | 156 | 1662 | 1431 |
| % in genes | | 1.6 | 12.18 | 2.77 | 1.82 |

| | | | | | |
|---|---|---|---|---|---|
| % in transposons | | 26 | 14.1 | 13.84 | 11.04 |
| % in repeats | | 5.36 | 4.49 | 7.04 | 5.59 |
| % in unclassified contigs | | 67.05 | 69.23 | 76.35 | 81.55 |
| >> within mixture AS | 1003 | - | - | 6682 | - |
| % in genes | 5.48 | | | 3.01 | |
| % in transposons | 20.34 | | | 14.65 | |
| % in repeats | 4.19 | | | 6.84 | |
| % in unclassified contigs | 69.99 | | | 75.5 | |
| AS: mixture vs. monoculture | - | - | - | 365 | - |
| % in genes | | | | 2.47 | |
| % in transposons | | | | 18.9 | |
| % in repeats | | | | 7.67 | |
| % in unclassified contigs | | | | 70.96 | |
| >> within supp2014 SH | - | - | - | 721 | - |
| % in genes | | | | 2.64 | |
| % in transposons | | | | 19.56 | |
| % in repeats | | | | 9.29 | |
| % in unclassified contigs | | | | 68.52 | |
| >> within monoculture SH | 546 | 171 | 164 | 1162 | 732 |
| % in genes | 3.85 | 2.92 | 20.12 | 2.07 | 1.5 |
| % in transposons | 24.36 | 29.82 | 11.59 | 18.42 | 21.17 |
| % in repeats | 5.31 | 7.02 | 10.37 | 7.4 | 3.14 |
| % in unclassified contigs | 66.48 | 60.23 | 57.93 | 72.12 | 74.18 |
| >> within mixed culture SH | 3118 | 519 | 310 | 825 | 800 |
| % in genes | 4.55 | 0.96 | 3.87 | 3.15 | 2.62 |
| % in transposons | 15.43 | 24.47 | 11.94 | 21.09 | 15.75 |
| % in repeats | 5.45 | 4.43 | 6.13 | 6.18 | 4.25 |
| % in unclassified contigs | 74.57 | 70.13 | 78.06 | 69.58 | 77.38 |
| Total tested | 753839 | 428603 | 134020 | 1445516 | 728159 |

**Table S11:** Number and percentages of cytosines with significant differences (FDR < 0.01) in DNA methylation between supp2002 plants and the other selection histories (supp2014, monoculture, mixture) in all sequence context or separate within the CG, CHG, and CHH contexts.

| Comparison to supp2002 | ALL | CG | CHG | CHH |
|---|---|---|---|---|
| supp2014 (monoculture assembly) | 21116 (2.00%) | 3664 (2.30%) | 6073 (3.58%) | 11379 (1.56%) |
| % in genes | 5.47 | 14.41 | 5.35 | 2.65 |
| % in transposons | 15.9 | 8.3 | 16.43 | 18.06 |
| % in repeats | 4.49 | 3.71 | 3.23 | 5.42 |
| % in unclassified contigs | 74.14 | 73.58 | 74.99 | 73.87 |
| monoculture history (both assemblies) | 44117 (4.17%) | 8610 (5.41%) | 12904 (7.61%) | 22603 (3.10%) |
| % in genes | 6.59 | 16.88 | 5.87 | 3.08 |
| % in transposons | 18.78 | 9.19 | 20.92 | 21.2 |
| % in repeats | 3.84 | 3.61 | 3.19 | 4.3 |
| % in unclassified contigs | 70.8 | 70.33 | 70.02 | 71.42 |
| >> only monoculture assembly | 25831 (2.44%) | 4856 (3.05%) | 7345 (4.33%) | 13630 (1.87%) |
| % in genes | 5.91 | 15.59 | 5.49 | 2.69 |
| % in transposons | 17.14 | 8.88 | 20.44 | 18.31 |
| % in repeats | 4.35 | 3.89 | 3.5 | 4.97 |
| % in unclassified contigs | 72.6 | 71.64 | 70.58 | 74.03 |
| >> only mixture assembly | 24637 (2.33%) | 5035 (3.05%) | 8120 (4.79%) | 11482 (1.58%) |
| % in genes | 6.26 | 15.25 | 5.21 | 3.06 |
| % in transposons | 17.82 | 9.02 | 19.79 | 20.29 |
| % in repeats | 4.08 | 3.61 | 3.29 | 4.83 |
| % in unclassified contigs | 71.84 | 72.12 | 71.71 | 71.82 |
| mixture history (both assemblies) | 26749 (2.53%) | 4937 (3.10%) | 8267 (4.88%) | 13545 (1.86%) |
| % in genes | 6.1 | 15.64 | 5.2 | 3.17 |
| % in transposons | 17.97 | 8.59 | 20.59 | 19.79 |
| % in repeats | 4.26 | 3.67 | 3.76 | 4.78 |
| % in unclassified contigs | 71.67 | 72.11 | 70.45 | 72.26 |
| >> only monoculture assembly | 28286 (2.68%) | 4560 (2.86%) | 9147 (5.40%) | 14579 (2.00%) |
| % in genes | 5.54 | 15.86 | 4.66 | 2.86 |
| % in transposons | 18.49 | 9.76 | 21.8 | 19.15 |
| % in repeats | 4.07 | 3.33 | 3.45 | 4.68 |
| % in unclassified contigs | 71.9 | 71.05 | 70.09 | 73.3 |
| >> only mixture assembly | 8702 (0.82%) | 2237 (1.40%) | 2996 (1.77%) | 3469 (0.48%) |
| % in genes | 7.22 | 14.26 | 5.71 | 3.98 |
| % in transposons | 14.28 | 7.82 | 16.02 | 16.95 |
| % in repeats | 4.25 | 3.17 | 3.81 | 5.33 |
| % in unclassified contigs | 74.25 | 74.74 | 74.47 | 73.74 |
| Total (percentage DMCs of tested cytosines) | 78241 (7.40%) | 16033 (10.07%) | 24001 (14.16%) | 38207 (5.25%) |
| Total cytosines tested | 1056852 | 159223 | 169470 | 728159 |

**Table S12:** Number of reads per sample and sample annotation. Samples which were removed are marked as "REMOVED".

| Sample ID | History | Assembly | Species | Number of reads |
|---|---|---|---|---|
| **Gal_mol_61** | REMOVED | REMOVED | *G. mollugo* | 1125191 |
| **Gal_mol_62** | mono | mixture | *G. mollugo* | 3160038 |
| **Gal_mol_63** | mono | mixture | *G. mollugo* | 2626973 |
| **Gal_mol_64** | mix | monoculture | *G. mollugo* | 1087619 |
| **Gal_mol_65** | mono | monoculture | *G. mollugo* | 2907198 |
| **Gal_mol_66** | mono | monoculture | *G. mollugo* | 1590521 |
| **Gal_mol_67** | mix | mixture | *G. mollugo* | 2939351 |
| **Gal_mol_68** | mix | monoculture | *G. mollugo* | 2042303 |
| **Gal_mol_69** | mix | monoculture | *G. mollugo* | 2345770 |
| **Gal_mol_70** | mix | mixture | *G. mollugo* | 1350313 |
| **Gal_mol_71** | mono | mixture | *G. mollugo* | 1936927 |
| **Gal_mol_72** | mono | mixture | *G. mollugo* | 2128368 |
| **Lat_pra_55** | mono | mixture | *L. pratensis* | 2148498 |
| **Lat_pra_56** | mix | monoculture | *L. pratensis* | 4084256 |
| **Lat_pra_57** | mono | monoculture | *L. pratensis* | 2927194 |
| **Lat_pra_58** | REMOVED | REMOVED | *L. pratensis* | 1347733 |
| **Lat_pra_59** | none | monoculture | *L. pratensis* | 1861493 |
| **Lat_pra_60** | none | monoculture | *L. pratensis* | 1903585 |
| **Lat_pra_61** | none | monoculture | *L. pratensis* | 3020936 |
| **Lat_pra_62** | mix | monoculture | *L. pratensis* | 2514618 |
| **Lat_pra_63** | mono | mixture | *L. pratensis* | 2470261 |
| **Lat_pra_64** | REMOVED | REMOVED | *L. pratensis* | 1718642 |
| **Lat_pra_65** | mono | monoculture | *L. pratensis* | 2035910 |
| **Lat_pra_66** | mix | monoculture | *L. pratensis* | 2657584 |
| **Pla_lan_61** | none | monoculture | *P. lanceolata* | 1030303 |
| **Pru_vul_79** | REMOVED | REMOVED | *P. vulgaris* | 3309366 |
| **Pla_lan_62** | mix | monoculture | *P. lanceolata* | 1937705 |
| **Pla_lan_63** | mono | mixture | *P. lanceolata* | 951736 |
| **Pla_lan_64** | mix | monoculture | *P. lanceolata* | 2643734 |
| **Pla_lan_65** | mono | mixture | *P. lanceolata* | 1490415 |
| **Pla_lan_66** | mix | mixture | *P. lanceolata* | 2964078 |
| **Pla_lan_67** | none | monoculture | *P. lanceolata* | 2666877 |
| **Pla_lan_68** | mono | mixture | *P. lanceolata* | 1187969 |
| **Pla_lan_69** | mono | monoculture | *P. lanceolata* | 1271285 |
| **Pru_vul_80** | REMOVED | REMOVED | *P. vulgaris* | 2117169 |
| **Pla_lan_70** | mix | mixture | *P. lanceolata* | 4236022 |

| | | | | |
|---|---|---|---|---|
| **Pla_lan_71** | mix | mixture | *P. lanceolata* | 2184886 |
| **Pla_lan_72** | mix | mixture | *P. lanceolata* | 3869524 |
| **Pla_lan_73** | mono | mixture | *P. lanceolata* | 3001306 |
| **Pla_lan_74** | mix | monoculture | *P. lanceolata* | 650039 |
| **Pla_lan_75** | mix | mixture | *P. lanceolata* | 3430054 |
| **Pla_lan_76** | mix | mixture | *P. lanceolata* | 1928321 |
| **Pla_lan_77** | none | monoculture | *P. lanceolata* | 3346471 |
| **Pla_lan_78** | mono | mixture | *P. lanceolata* | 2895468 |
| **Pla_lan_79** | mix | mixture | *P. lanceolata* | 2286933 |
| **Pla_lan_80** | mix | mixture | *P. lanceolata* | 738251 |
| **Pla_lan_81** | REMOVED | REMOVED | *P. lanceolata* | 310715 |
| **Pla_lan_82** | REMOVED | REMOVED | *P. lanceolata* | 2862864 |
| **Pru_vul_81** | mix | mixture | *P. vulgaris* | 717538 |
| **Pru_vul_82** | mix | monoculture | *P. vulgaris* | 3561439 |
| **Pru_vul_83** | mix | mixture | *P. vulgaris* | 2760156 |
| **Pru_vul_84** | mix | monoculture | *P. vulgaris* | 1169261 |
| **Pru_vul_85** | mono | monoculture | *P. vulgaris* | 3109074 |
| **Pru_vul_86** | none | monoculture | *P. vulgaris* | 1735603 |
| **Pru_vul_87** | REMOVED | REMOVED | *P. vulgaris* | 413191 |
| **Pru_vul_88** | none | mixture | *P. vulgaris* | 2177980 |
| **Pru_vul_89** | mix | mixture | *P. vulgaris* | 2156781 |
| **Pru_vul_90** | mix | monoculture | *P. vulgaris* | 1527449 |
| **Pru_vul_91** | mix | mixture | *P. vulgaris* | 1864795 |
| **Pru_vul_92** | mono | mixture | *P. vulgaris* | 2560557 |
| **Pru_vul_93** | mix | monoculture | *P. vulgaris* | 1419878 |
| **Pru_vul_94** | mono | mixture | *P. vulgaris* | 2160906 |
| **Pru_vul_95** | none | monoculture | *P. vulgaris* | 1832766 |
| **Pru_vul_96** | none | monoculture | *P. vulgaris* | 913030 |
| **Pru_vul_97** | mono | monoculture | *P. vulgaris* | 1436599 |
| **Pru_vul_98** | none | monoculture | *P. vulgaris* | 1173822 |
| **Pru_vul_99** | mono | mixture | *P. vulgaris* | 2028428 |
| **Pru_vul_100** | none | monoculture | *P. vulgaris* | 1623811 |
| **Ver_cha_73** | REMOVED | REMOVED | *V. chamaedrys* | 1728742 |
| **Pru_vul_101** | mono | monoculture | *P. vulgaris* | 1534763 |
| **Pru_vul_102** | mix | mixture | *P. vulgaris* | 1449176 |
| **Pru_vul_103** | none | monoculture | *P. vulgaris* | 1965743 |
| **Ver_cha_74** | mix | mixture | *V. chamaedrys* | 1591765 |
| **Ver_cha_75** | mix | mixture | *V. chamaedrys* | 2918977 |
| **Ver_cha_76** | mix | mixture | *V. chamaedrys* | 2528710 |

| | | | | |
|---|---|---|---|---|
| **Ver_cha_77** | none | monoculture | *V. chamaedrys* | 948161 |
| **Ver_cha_78** | mix | monoculture | *V. chamaedrys* | 3275173 |
| **Ver_cha_79** | mix | monoculture | *V. chamaedrys* | 1513726 |
| **Ver_cha_80** | none | monoculture | *V. chamaedrys* | 3064784 |
| **Ver_cha_81** | mono | monoculture | *V. chamaedrys* | 2273744 |
| **Ver_cha_82** | mono | monoculture | *V. chamaedrys* | 1039270 |
| **Ver_cha_83** | none | monoculture | *V. chamaedrys* | 1591189 |
| **Ver_cha_84** | none | monoculture | *V. chamaedrys* | 1549011 |
| **Ver_cha_85** | none | monoculture | *V. chamaedrys* | 2151267 |
| **Gal_mol_37** | REMOVED | REMOVED | *G. mollugo* | 803303 |
| **Gal_mol_38** | mix | mixture | *G. mollugo* | 2402333 |
| **Gal_mol_39** | mono | mixture | *G. mollugo* | 1974015 |
| **Gal_mol_40** | mono | mixture | *G. mollugo* | 808096 |
| **Gal_mol_41** | none | mixture | *G. mollugo* | 2621320 |
| **Gal_mol_42** | mix | monoculture | *G. mollugo* | 1638542 |
| **Gal_mol_43** | none | mixture | *G. mollugo* | 2148127 |
| **Gal_mol_44** | mono | mixture | *G. mollugo* | 1959705 |
| **Gal_mol_45** | mono | monoculture | *G. mollugo* | 2014941 |
| **Gal_mol_46** | mono | mixture | *G. mollugo* | 1231745 |
| **Gal_mol_47** | mix | mixture | *G. mollugo* | 1161455 |
| **Gal_mol_48** | mono | mixture | *G. mollugo* | 1286234 |
| **Gal_mol_49** | mono | mixture | *G. mollugo* | 1392987 |
| **Gal_mol_50** | mono | mixture | *G. mollugo* | 2355224 |
| **Gal_mol_51** | mono | monoculture | *G. mollugo* | 2205662 |
| **Gal_mol_52** | mix | mixture | *G. mollugo* | 944319 |
| **Gal_mol_53** | mix | mixture | *G. mollugo* | 2126141 |
| **Gal_mol_54** | mix | monoculture | *G. mollugo* | 1073676 |
| **Gal_mol_55** | none | mixture | *G. mollugo* | 1490760 |
| **Gal_mol_56** | mix | mixture | *G. mollugo* | 1115602 |
| **Gal_mol_57** | mix | mixture | *G. mollugo* | 1000609 |
| **Gal_mol_58** | mono | monoculture | *G. mollugo* | 1223323 |
| **Gal_mol_59** | mix | mixture | *G. mollugo* | 1161741 |
| **Gal_mol_60** | mix | mixture | *G. mollugo* | 1577366 |
| **Lat_pra_43** | mix | mixture | *L. pratensis* | 1792723 |
| **Lat_pra_44** | none | monoculture | *L. pratensis* | 2649804 |
| **Lat_pra_45** | mono | mixture | *L. pratensis* | 2260990 |
| **Lat_pra_46** | REMOVED | REMOVED | *L. pratensis* | 853947 |
| **Lat_pra_47** | none | monoculture | *L. pratensis* | 2151801 |
| **Lat_pra_48** | REMOVED | REMOVED | *L. pratensis* | 1538876 |

| | | | | |
|---|---|---|---|---|
| **Lat_pra_49** | mono | mixture | *L. pratensis* | 2426689 |
| **Lat_pra_50** | mix | monoculture | *L. pratensis* | 1829011 |
| **Lat_pra_51** | none | monoculture | *L. pratensis* | 1766347 |
| **Lat_pra_52** | none | monoculture | *L. pratensis* | 1290616 |
| **Lat_pra_53** | mix | mixture | *L. pratensis* | 1525508 |
| **Lat_pra_54** | none | monoculture | *L. pratensis* | 1498117 |
| **Pla_lan_49** | mix | mixture | *P. lanceolata* | 1104144 |
| **Pla_lan_50** | REMOVED | REMOVED | *P. lanceolata* | 3098751 |
| **Pla_lan_51** | none | monoculture | *P. lanceolata* | 2106272 |
| **Pla_lan_52** | REMOVED | REMOVED | *P. lanceolata* | 460198 |
| **Pla_lan_53** | mono | mixture | *P. lanceolata* | 1034110 |
| **Pla_lan_54** | mono | monoculture | *P. lanceolata* | 1010179 |
| **Pla_lan_55** | mono | mixture | *P. lanceolata* | 1916689 |
| **Pla_lan_56** | none | monoculture | *P. lanceolata* | 1752511 |
| **Pla_lan_57** | mix | mixture | *P. lanceolata* | 2123750 |
| **Pla_lan_58** | REMOVED | REMOVED | *P. lanceolata* | 1186507 |
| **Pla_lan_59** | none | monoculture | *P. lanceolata* | 2119342 |
| **Pla_lan_60** | mono | mixture | *P. lanceolata* | 1432527 |
| **Ver_cha_61** | none | monoculture | *V. chamaedrys* | 1314927 |
| **Ver_cha_62** | mix | mixture | *V. chamaedrys* | 3433475 |
| **Ver_cha_63** | none | monoculture | *V. chamaedrys* | 2902452 |
| **Ver_cha_64** | mono | mixture | *V. chamaedrys* | 1562507 |
| **Ver_cha_65** | mix | monoculture | *V. chamaedrys* | 3824333 |
| **Ver_cha_66** | none | monoculture | *V. chamaedrys* | 1909073 |
| **Ver_cha_67** | mix | mixture | *V. chamaedrys* | 4126881 |
| **Ver_cha_68** | mono | mixture | *V. chamaedrys* | 3747489 |
| **Ver_cha_69** | none | monoculture | *V. chamaedrys* | 3156590 |
| **Ver_cha_70** | mix | mixture | *V. chamaedrys* | 1749794 |
| **Ver_cha_71** | mono | monoculture | *V. chamaedrys* | 3123304 |
| **Ver_cha_72** | mono | mixture | *V. chamaedrys* | 2578881 |
| **Pru_vul_55** | mono | mixture | *P. vulgaris* | 691470 |
| **Pru_vul_56** | none | mixture | *P. vulgaris* | 2739578 |
| **Pru_vul_57** | mono | mixture | *P. vulgaris* | 2596880 |
| **Pru_vul_58** | mono | mixture | *P. vulgaris* | 1168800 |
| **Pru_vul_59** | mono | monoculture | *P. vulgaris* | 2898503 |
| **Pru_vul_60** | REMOVED | REMOVED | *P. vulgaris* | 355738 |
| **Pru_vul_61** | mix | mixture | *P. vulgaris* | 2625754 |
| **Pru_vul_62** | mix | mixture | *P. vulgaris* | 2739524 |
| **Pru_vul_63** | none | monoculture | *P. vulgaris* | 2605891 |

| | | | | |
|---|---|---|---|---|
| **Pru_vul_64** | none | mixture | *P. vulgaris* | 1431419 |
| **Pru_vul_65** | none | monoculture | *P. vulgaris* | 2101842 |
| **Pru_vul_66** | mix | mixture | *P. vulgaris* | 1588973 |
| **Pru_vul_67** | mono | mixture | *P. vulgaris* | 1490269 |
| **Pru_vul_68** | mono | mixture | *P. vulgaris* | 5590228 |
| **Pru_vul_69** | mono | mixture | *P. vulgaris* | 3852766 |
| **Pru_vul_70** | mix | mixture | *P. vulgaris* | 1384796 |
| **Pru_vul_71** | mix | mixture | *P. vulgaris* | 3634177 |
| **Pru_vul_72** | none | monoculture | *P. vulgaris* | 1536592 |
| **Pru_vul_73** | mix | monoculture | *P. vulgaris* | 2969939 |
| **Pru_vul_74** | mix | mixture | *P. vulgaris* | 2588639 |
| **Pru_vul_75** | mix | mixture | *P. vulgaris* | 1945543 |
| **Pru_vul_76** | none | monoculture | *P. vulgaris* | 1916058 |
| **Pru_vul_77** | mono | mixture | *P. vulgaris* | 2257263 |
| **Pru_vul_78** | mono | mixture | *P. vulgaris* | 2636368 |
| **Gal_mol_73** | mono | monoculture | *G. mollugo* | 1163227 |
| **Gal_mol_74** | mix | mixture | *G. mollugo* | 3257717 |
| **Gal_mol_75** | mix | mixture | *G. mollugo* | 1628121 |
| **Gal_mol_76** | mono | mixture | *G. mollugo* | 971915 |
| **Gal_mol_77** | mix | mixture | *G. mollugo* | 1581366 |
| **Gal_mol_78** | none | mixture | *G. mollugo* | 1225253 |
| **Gal_mol_79** | mix | monoculture | *G. mollugo* | 1677836 |
| **Gal_mol_80** | mono | mixture | *G. mollugo* | 1584435 |
| **Gal_mol_81** | mix | mixture | *G. mollugo* | 1115013 |
| **Gal_mol_82** | mono | mixture | *G. mollugo* | 1117064 |
| **Gal_mol_83** | mix | mixture | *G. mollugo* | 976028 |
| **Gal_mol_84** | none | mixture | *G. mollugo* | 1636130 |
| **Gal_mol_85** | mix | mixture | *G. mollugo* | 1663543 |
| **Gal_mol_86** | mix | mixture | *G. mollugo* | 3234733 |
| **Gal_mol_87** | mix | mixture | *G. mollugo* | 2511044 |
| **Gal_mol_88** | none | mixture | *G. mollugo* | 1002156 |
| **Lat_pra_67** | mono | monoculture | *L. pratensis* | 3688530 |
| **Lat_pra_68** | mix | mixture | *L. pratensis* | 2248028 |
| **Lat_pra_69** | mix | mixture | *L. pratensis* | 3179381 |
| **Lat_pra_70** | mono | mixture | *L. pratensis* | 2175697 |
| **Lat_pra_71** | mix | monoculture | *L. pratensis* | 2583851 |
| **Lat_pra_72** | none | monoculture | *L. pratensis* | 1977899 |
| **Lat_pra_73** | mix | monoculture | *L. pratensis* | 2521592 |
| **Lat_pra_74** | mono | monoculture | *L. pratensis* | 2880947 |

| | | | | |
|---|---|---|---|---|
| **Lat_pra_75** | mono | monoculture | *L. pratensis* | 2645085 |
| **Lat_pra_76** | none | monoculture | *L. pratensis* | 4566757 |
| **Lat_pra_77** | none | monoculture | *L. pratensis* | 3696714 |
| **Pla_lan_83** | REMOVED | REMOVED | *P. lanceolata* | 285465 |
| **Pla_lan_84** | mix | monoculture | *P. lanceolata* | 1318074 |
| **Pla_lan_85** | REMOVED | REMOVED | *P. lanceolata* | 728091 |
| **Pla_lan_86** | mix | mixture | *P. lanceolata* | 1549355 |
| **Pla_lan_87** | mono | monoculture | *P. lanceolata* | 1106823 |
| **Pla_lan_88** | mix | monoculture | *P. lanceolata* | 1108928 |
| **Pla_lan_89** | mix | mixture | *P. lanceolata* | 726230 |
| **Pla_lan_90** | none | monoculture | *P. lanceolata* | 643494 |
| **Pla_lan_91** | mix | mixture | *P. lanceolata* | 625937 |
| **Pla_lan_92** | REMOVED | REMOVED | *P. lanceolata* | 796798 |
| **Pla_lan_93** | mono | mixture | *P. lanceolata* | 2391541 |
| **Pla_lan_94** | mono | mixture | *P. lanceolata* | 1522609 |
| **Pla_lan_95** | mix | mixture | *P. lanceolata* | 527985 |
| **Pla_lan_96** | mix | monoculture | *P. lanceolata* | 1514123 |
| **Pla_lan_97** | mix | mixture | *P. lanceolata* | 591931 |
| **Pla_lan_98** | mix | mixture | *P. lanceolata* | 2014206 |
| **Pla_lan_99** | mono | mixture | *P. lanceolata* | 801859 |
| **Pla_lan_100** | mono | mixture | *P. lanceolata* | 772504 |
| **Pla_lan_101** | mono | mixture | *P. lanceolata* | 798581 |
| **Pla_lan_102** | mono | monoculture | *P. lanceolata* | 1050566 |
| **Pla_lan_103** | none | monoculture | *P. lanceolata* | 1483012 |
| **Pla_lan_104** | mono | monoculture | *P. lanceolata* | 870786 |
| **Pla_lan_105** | mono | mixture | *P. lanceolata* | 2266757 |
| **Pla_lan_106** | mono | mixture | *P. lanceolata* | 936675 |
| **Pla_lan_107** | mono | mixture | *P. lanceolata* | 809294 |
| **Pla_lan_108** | mix | mixture | *P. lanceolata* | 1630091 |
| **Pla_lan_109** | mix | mixture | *P. lanceolata* | 1563374 |
| **Pla_lan_110** | mix | mixture | *P. lanceolata* | 2781459 |
| **Pla_lan_111** | REMOVED | REMOVED | *P. lanceolata* | 2810543 |
| **Pla_lan_112** | mix | mixture | *P. lanceolata* | 1867924 |
| **Pla_lan_113** | mix | mixture | *P. lanceolata* | 1457906 |
| **Pru_vul_104** | none | mixture | *P. vulgaris* | 1221866 |
| **Pru_vul_105** | none | mixture | *P. vulgaris* | 1805129 |
| **Pru_vul_106** | mono | mixture | *P. vulgaris* | 1500719 |
| **Pru_vul_107** | mix | mixture | *P. vulgaris* | 3737284 |
| **Pru_vul_108** | REMOVED | REMOVED | *P. vulgaris* | 4565778 |

| | | | | |
|---|---|---|---|---|
| **Pru_vul_109** | mono | monoculture | *P. vulgaris* | 2333415 |
| **Pru_vul_110** | mono | mixture | *P. vulgaris* | 4117321 |
| **Pru_vul_111** | mono | monoculture | *P. vulgaris* | 2703139 |
| **Pru_vul_112** | mono | mixture | *P. vulgaris* | 3055922 |
| **Pru_vul_113** | mono | mixture | *P. vulgaris* | 2846477 |
| **Pru_vul_114** | mono | mixture | *P. vulgaris* | 2133140 |
| **Pru_vul_115** | mix | mixture | *P. vulgaris* | 2930887 |
| **Pru_vul_116** | mono | mixture | *P. vulgaris* | 3784676 |
| **Pru_vul_117** | mix | mixture | *P. vulgaris* | 4127007 |
| **Pru_vul_118** | mix | monoculture | *P. vulgaris* | 3165061 |
| **Pru_vul_119** | REMOVED | REMOVED | *P. vulgaris* | 5679189 |
| **Pru_vul_120** | mono | mixture | *P. vulgaris* | 4566982 |
| **Pru_vul_121** | mix | mixture | *P. vulgaris* | 2595625 |
| **Pru_vul_122** | REMOVED | REMOVED | *P. vulgaris* | 5789083 |
| **Ver_cha_86** | none | monoculture | *V. chamaedrys* | 2406739 |
| **Ver_cha_87** | mix | monoculture | *V. chamaedrys* | 3427303 |
| **Ver_cha_88** | mix | monoculture | *V. chamaedrys* | 3021857 |
| **Ver_cha_89** | mono | monoculture | *V. chamaedrys* | 2730216 |
| **Ver_cha_90** | mono | mixture | *V. chamaedrys* | 2537468 |
| **Ver_cha_91** | mono | monoculture | *V. chamaedrys* | 3142096 |
| **Ver_cha_92** | mix | monoculture | *V. chamaedrys* | 3099042 |
| **Ver_cha_93** | mono | mixture | *V. chamaedrys* | 2217280 |
| **Ver_cha_94** | REMOVED | REMOVED | *V. chamaedrys* | 4169075 |
| **Ver_cha_95** | mono | mixture | *V. chamaedrys* | 3659100 |
| **Ver_cha_96** | mono | monoculture | *V. chamaedrys* | 1466806 |
| **Ver_cha_97** | mono | mixture | *V. chamaedrys* | 3803337 |
| **Gal_mol_1** | mix | monoculture | *G. mollugo* | 3077771 |
| **Gal_mol_2** | mix | monoculture | *G. mollugo* | 3270499 |
| **Gal_mol_3** | mix | monoculture | *G. mollugo* | 2783616 |
| **Gal_mol_4** | REMOVED | REMOVED | *G. mollugo* | 716864 |
| **Gal_mol_5** | mix | monoculture | *G. mollugo* | 4943175 |
| **Gal_mol_6** | mix | monoculture | *G. mollugo* | 2166392 |
| **Gal_mol_31** | mono | monoculture | *G. mollugo* | 4194097 |
| **Gal_mol_32** | mono | monoculture | *G. mollugo* | 2458479 |
| **Gal_mol_33** | mono | monoculture | *G. mollugo* | 3259615 |
| **Gal_mol_34** | mono | monoculture | *G. mollugo* | 1957792 |
| **Gal_mol_35** | mono | monoculture | *G. mollugo* | 2209035 |
| **Gal_mol_36** | mono | monoculture | *G. mollugo* | 1976646 |
| **Lat_pra_7** | mix | monoculture | *L. pratensis* | 2546409 |

| | | | | |
|---|---|---|---|---|
| **Lat_pra_8** | mix | monoculture | *L. pratensis* | 4502883 |
| **Lat_pra_9** | mix | monoculture | *L. pratensis* | 3005350 |
| **Lat_pra_10** | REMOVED | REMOVED | *L. pratensis* | 1451398 |
| **Lat_pra_11** | mix | monoculture | *L. pratensis* | 2767910 |
| **Lat_pra_12** | mix | monoculture | *L. pratensis* | 2511875 |
| **Lat_pra_37** | mono | monoculture | *L. pratensis* | 3456836 |
| **Lat_pra_38** | mono | monoculture | *L. pratensis* | 2694232 |
| **Lat_pra_39** | mono | monoculture | *L. pratensis* | 3538764 |
| **Lat_pra_40** | mono | monoculture | *L. pratensis* | 3048057 |
| **Lat_pra_41** | mono | monoculture | *L. pratensis* | 2571595 |
| **Lat_pra_42** | mono | monoculture | *L. pratensis* | 3208389 |
| **Pla_lan_13** | mix | monoculture | *P. lanceolata* | 2279166 |
| **Pla_lan_14** | mix | monoculture | *P. lanceolata* | 2944789 |
| **Pla_lan_15** | REMOVED | REMOVED | *P. lanceolata* | 784330 |
| **Pla_lan_16** | REMOVED | REMOVED | *P. lanceolata* | 863073 |
| **Pla_lan_17** | mix | monoculture | *P. lanceolata* | 2242254 |
| **Pla_lan_18** | mix | monoculture | *P. lanceolata* | 1598081 |
| **Pla_lan_43** | mono | monoculture | *P. lanceolata* | 1287372 |
| **Pla_lan_44** | mono | monoculture | *P. lanceolata* | 919800 |
| **Pla_lan_45** | mono | monoculture | *P. lanceolata* | 1498272 |
| **Pla_lan_46** | mono | monoculture | *P. lanceolata* | 802658 |
| **Pla_lan_47** | mono | monoculture | *P. lanceolata* | 1828560 |
| **Pla_lan_48** | mono | monoculture | *P. lanceolata* | 1484134 |
| **Pru_vul_19** | mix | monoculture | *P. vulgaris* | 1271540 |
| **Pru_vul_20** | mix | monoculture | *P. vulgaris* | 3960379 |
| **Pru_vul_21** | mix | monoculture | *P. vulgaris* | 2275789 |
| **Pru_vul_22** | REMOVED | REMOVED | *P. vulgaris* | 906235 |
| **Pru_vul_23** | mix | monoculture | *P. vulgaris* | 1871631 |
| **Pru_vul_24** | REMOVED | REMOVED | *P. vulgaris* | 1541075 |
| **Pru_vul_49** | mono | monoculture | *P. vulgaris* | 3346252 |
| **Pru_vul_50** | mono | monoculture | *P. vulgaris* | 1294205 |
| **Pru_vul_51** | mono | monoculture | *P. vulgaris* | 2788968 |
| **Pru_vul_52** | mono | monoculture | *P. vulgaris* | 1835662 |
| **Pru_vul_53** | mono | monoculture | *P. vulgaris* | 2550351 |
| **Pru_vul_54** | mono | monoculture | *P. vulgaris* | 2069899 |
| **Ver_cha_25** | mix | monoculture | *V. chamaedrys* | 3120608 |
| **Ver_cha_26** | mix | monoculture | *V. chamaedrys* | 4050406 |
| **Ver_cha_27** | mix | monoculture | *V. chamaedrys* | 1110442 |
| **Ver_cha_28** | mix | monoculture | *V. chamaedrys* | 1591843 |

| | | | | |
|---|---|---|---|---|
| **Ver_cha_29** | mix | monoculture | *V. chamaedrys* | 3418320 |
| **Ver_cha_30** | mix | monoculture | *V. chamaedrys* | 1692484 |
| **Ver_cha_55** | mono | monoculture | *V. chamaedrys* | 2897001 |
| **Ver_cha_56** | mono | monoculture | *V. chamaedrys* | 3381267 |
| **Ver_cha_57** | mono | monoculture | *V. chamaedrys* | 2312164 |
| **Ver_cha_58** | mono | monoculture | *V. chamaedrys* | 1379652 |
| **Ver_cha_59** | mono | monoculture | *V. chamaedrys* | 1567552 |
| **Ver_cha_60** | mono | monoculture | *V. chamaedrys* | 1738241 |
| **Veronica_1** | founder | ukn | *V. chamaedrys* | 365521 |
| **Veronica_2** | founder | ukn | *V. chamaedrys* | 2063137 |
| **Veronica_3** | founder | ukn | *V. chamaedrys* | 1717447 |
| **Veronica_4** | founder | ukn | *V. chamaedrys* | 670181 |
| **Veronica_5** | founder | ukn | *V. chamaedrys* | 1539973 |
| **Veronica_6** | founder | ukn | *V. chamaedrys* | 1290497 |
| **Veronica_7** | founder | ukn | *V. chamaedrys* | 1491379 |
| **Veronica_8** | founder | ukn | *V. chamaedrys* | 1279083 |
| **Veronica_9** | founder | ukn | *V. chamaedrys* | 1070581 |
| **Veronica_10** | founder | ukn | *V. chamaedrys* | 740728 |
| **Veronica_11** | founder | ukn | *V. chamaedrys* | 873518 |
| **Veronica_12** | founder | ukn | *V. chamaedrys* | 1436517 |
| **Veronica_13** | founder | ukn | *V. chamaedrys* | 397524 |
| **Veronica_14** | founder | ukn | *V. chamaedrys* | 1896753 |
| **Veronica_15** | founder | ukn | *V. chamaedrys* | 1675914 |
| **Veronica_16** | founder | ukn | *V. chamaedrys* | 648840 |
| **Veronica_17** | founder | ukn | *V. chamaedrys* | 1373855 |
| **Veronica_18** | founder | ukn | *V. chamaedrys* | 1270269 |
| **Veronica_19** | founder | ukn | *V. chamaedrys* | 1637242 |
| **Veronica_20** | founder | ukn | *V. chamaedrys* | 1205803 |
| **Veronica_21** | founder | ukn | *V. chamaedrys* | 968156 |
| **Veronica_22** | founder | ukn | *V. chamaedrys* | 638912 |
| **Veronica_23** | founder | ukn | *V. chamaedrys* | 800122 |
| **Veronica_24** | founder | ukn | *V. chamaedrys* | 1183780 |
| **Veronica_25** | founder | ukn | *V. chamaedrys* | 945155 |
| **Veronica_26** | founder | ukn | *V. chamaedrys* | 2195802 |
| **Veronica_27** | founder | ukn | *V. chamaedrys* | 1933462 |
| **Veronica_28** | founder | ukn | *V. chamaedrys* | 479313 |
| **Veronica_29** | founder | ukn | *V. chamaedrys* | 1171403 |
| **Veronica_30** | founder | ukn | *V. chamaedrys* | 859996 |
| **Veronica_31** | founder | ukn | *V. chamaedrys* | 1296597 |

| | | | | |
|---|---|---|---|---|
| **Veronica_32** | founder | ukn | *V. chamaedrys* | 945812 |
| **Veronica_33** | founder | ukn | *V. chamaedrys* | 828486 |
| **Veronica_34** | founder | ukn | *V. chamaedrys* | 623108 |
| **Veronica_35** | founder | ukn | *V. chamaedrys* | 722030 |
| **Veronica_36** | founder | ukn | *V. chamaedrys* | 884857 |
| **Veronica_37** | founder | ukn | *V. chamaedrys* | 1982938 |
| **Veronica_38** | founder | ukn | *V. chamaedrys* | 3490690 |
| **Veronica_39** | founder | ukn | *V. chamaedrys* | 3201218 |
| **Veronica_40** | founder | ukn | *V. chamaedrys* | 1061100 |
| **Veronica_54** | founder | singleSeedPot | *V. chamaedrys* | 966290 |
| **Veronica_55** | founder | singleSeedPot | *V. chamaedrys* | 1613695 |
| **Veronica_56** | founder | singleSeedPot | *V. chamaedrys* | 1273192 |
| **Veronica_57** | REMOVED | REMOVED | *V. chamaedrys* | 45935 |
| **Veronica_58** | REMOVED | REMOVED | *V. chamaedrys* | 137503 |
| **Veronica_59** | founder | singleSeedPot | *V. chamaedrys* | 911688 |
| **Veronica_60** | founder | singleSeedPot | *V. chamaedrys* | 1179215 |

**Table S13:** Average number of reads per treatment combination.

| Species | Selection history | Assembly | Average | Standard deviation |
|---|---|---|---|---|
| ***G. mollugo*** | mixture | mixture | 1,763,711 | 789,012 |
| ***L. pratensis*** | mixture | mixture | 2,186,410 | 726,078 |
| ***P. lanceolata*** | mixture | mixture | 1,914,398 | 1,071,444 |
| ***P. vulgaris*** | mixture | mixture | 2,427,916 | 927,669 |
| ***V. chamaedrys*** | mixture | mixture | 2,724,934 | 977,611 |
| ***G. mollugo*** | monoculture | mixture | 1,752,409 | 675,250 |
| ***L. pratensis*** | monoculture | mixture | 2,296,427 | 145,689 |
| ***P. lanceolata*** | monoculture | mixture | 1,513,128 | 760,618 |
| ***P. vulgaris*** | monoculture | mixture | 2,724,398 | 1,261,751 |
| ***V. chamaedrys*** | monoculture | mixture | 2,872,295 | 875,086 |
| ***G. mollugo*** | supp2014 | mixture | 1,687,291 | 601,483 |
| ***P. vulgaris*** | supp2014 | mixture | 1,874,194 | 605,707 |
| ***G. mollugo*** | mixture | monoculture | 2,373,382 | 1,122,164 |
| ***L. pratensis*** | mixture | monoculture | 2,865,940 | 765,545 |
| ***P. lanceolata*** | mixture | monoculture | 1,823,689 | 716,140 |
| ***P. vulgaris*** | mixture | monoculture | 2,319,237 | 1,023,793 |
| ***V. chamaedrys*** | mixture | monoculture | 2,762,128 | 1,001,542 |
| ***G. mollugo*** | monoculture | monoculture | 2,263,378 | 861,348 |
| ***L. pratensis*** | monoculture | monoculture | 2,972,322 | 486,033 |
| ***P. lanceolata*** | monoculture | monoculture | 1,193,676 | 314,465 |
| ***P. vulgaris*** | monoculture | monoculture | 2,325,069 | 685,999 |
| ***V. chamaedrys*** | monoculture | monoculture | 2,254,276 | 800,913 |
| ***L. pratensis*** | supp2014 | monoculture | 2,398,552 | 1,002,301 |
| ***P. lanceolata*** | supp2014 | monoculture | 1,893,535 | 870,264 |
| ***P. vulgaris*** | supp2014 | monoculture | 1,740,516 | 474,996 |
| ***V. chamaedrys*** | supp2014 | monoculture | 2,099,419 | 769,548 |
| ***V. chamaedrys*** | supp2002 | single seed pod | 1,188,816 | 280,201 |

| | | | | |
|---|---|---|---|---|
| ***V. chamaedrys*** | supp2002 | garden | 1,271,442 | 675,447 |
